# Reconstructing the Molecular Life History of Gliomas

**DOI:** 10.1101/192369

**Authors:** Floris P. Barthel, Pieter Wesseling, Roel G.W. Verhaak

**Affiliations:** The Jackson Laboratory for Genomic Medicine, Farmington, CT 06030; Department of Pathology, VU University Medical Center/Brain Tumor Center Amsterdam, Amsterdam, The Netherlands.; Department of Pathology, Princess Máxima Center for Pediatric Oncology and University Medical Center Utrecht, Utrecht, The Netherlands.

**Author notes:** Correspondence should be addressed to F.P.B.

## Abstract

At the time of clinical presentation, the very heterogeneous group of pediatric and adult gliomas carry a wide range of diverse somatic genomic alterations. These include chromosome-sized gains and losses, focal amplification and deletions, rearrangements resulting in transcript fusions, small insertions/deletions, and point mutations. Tumor cells pay a penalty for maintaining these abnormalities which therefore must provide cells with a competitive advantage to become engrained into the glioma genome. Here, we propose a model for gliomagenesis consisting of five consecutive phases that glioma cells have traversed prior to diagnosis. Tumor growth is repressed by activated DNA damage response pathways and dysfunctional telomeres in physiological conditions. Disruption of the p16-RB-p53 pathway and the acquisition of a telomere maintenance mechanism can bypass these bottlenecks. We relate somatic alterations to each of these steps, in order to reconstruct the life history of glioma. Understanding the story that each glioma tells at presentation may facilitate the design of novel, more effective therapeutic approaches.

**Key Concepts:** *Glioma initiating event*: The first event that initiates the clonal expansion of cells

*Oncogene-induced senescence*: Durable growth arrest triggered by continued oncogene exposure

*Replicative senescence*: Durable growth arrest triggered via telomere dysfunction and activated DNA damage pathways

*Crisis*: Widespread cell death triggered via telomere dysfunction

*Senescence bypass event*: Any molecular alteration that bypasses or suppresses oncogene-induced senescence

*Senescence-associated secretory phenotype (SASP)*: Senescent cells secrete various immunogenic cytokines, growth factors and proteases into the microenvironment

*Functional redundancy*: Used to describe two or more genomic changes that provide overlapping functional effect

*Neutral evolution:* changes due to stochastic allelic variation that do not affect fitness

*Selective sweep:* The elimination of genetic variation following strong positive selection effectively reducing the tumor to a single clone

*Clonal event*: Somatic mutation or copy number event that is conserved across all tumor cells

*Subclonal event*: Somatic mutation or copy number event that is only present in a subset (subclone) of tumor cells

*Chromothripsis*: A punctuated shattering of genomic DNA

*Kataegis*: Clustered regions of hypermutation

*Polyploidization*: The multiplication of chromosome content in a cell

*Breakage fusion bridge (BFB) cycle*: Cyclic fusion of uncapped telomeres, bridge formation during anaphase and subsequent breakage leading to unequal inheritance of DNA

*Dicentric chromosome*: Two fused chromosomes span across the mitotic spindle in anaphase, called dicentric because it has two centromeres

*Double minute (DM) chromosome*: Extra-chromosomal circular DNA segment lacking centromere(s) and telomeres

*Immortalization event*: The last straw in the immortalization process that directly leads to telomere stabilization

## 1. Introduction

As with other human cancers, the pathogenesis and molecular evolution of glial tumors of the central nervous system (CNS) is often characterized by chromosomal aberrations, widespread or focal copy number changes and targeted gain and loss of function events in oncogenes and tumor suppressor genes [1-3]. Various permutations of somatic alterations are associated with distinct tumor entities and differential sensitivities to treatment, such as a chromosome 1p/19q-codeletion in oligodendrogliomas conferring increased sensitivity to chemotherapy [4, 5]. In this review, we propose five phases in gliomagenesis (Figure 1) that occur sequentially and ultimately lead to the tumor at clinical diagnosis. Each phase is characterized by distinct molecular alterations and phenotypic characteristics, such as differences in growth dynamics and evolutionary mechanisms. A critical assumption in our model is the existence of two growth barriers, which we refer to as oncogene-induced and replicative senescence. Similar barriers have been described in detail in the context of cultured epithelial cells and fibroblasts and much of this work has paved the road for our understanding of these mechanisms in gliomagenesis [6-8]. We systematically review the evidence for our model and propose candidate mechanisms where definitive evidence is lacking. We discuss in depth our model across various entities of diffuse glioma recognized by the most recent WHO classification and touch upon a few examples in the realm of the (much less frequent) non-diffuse gliomas.

**Figure 1.**
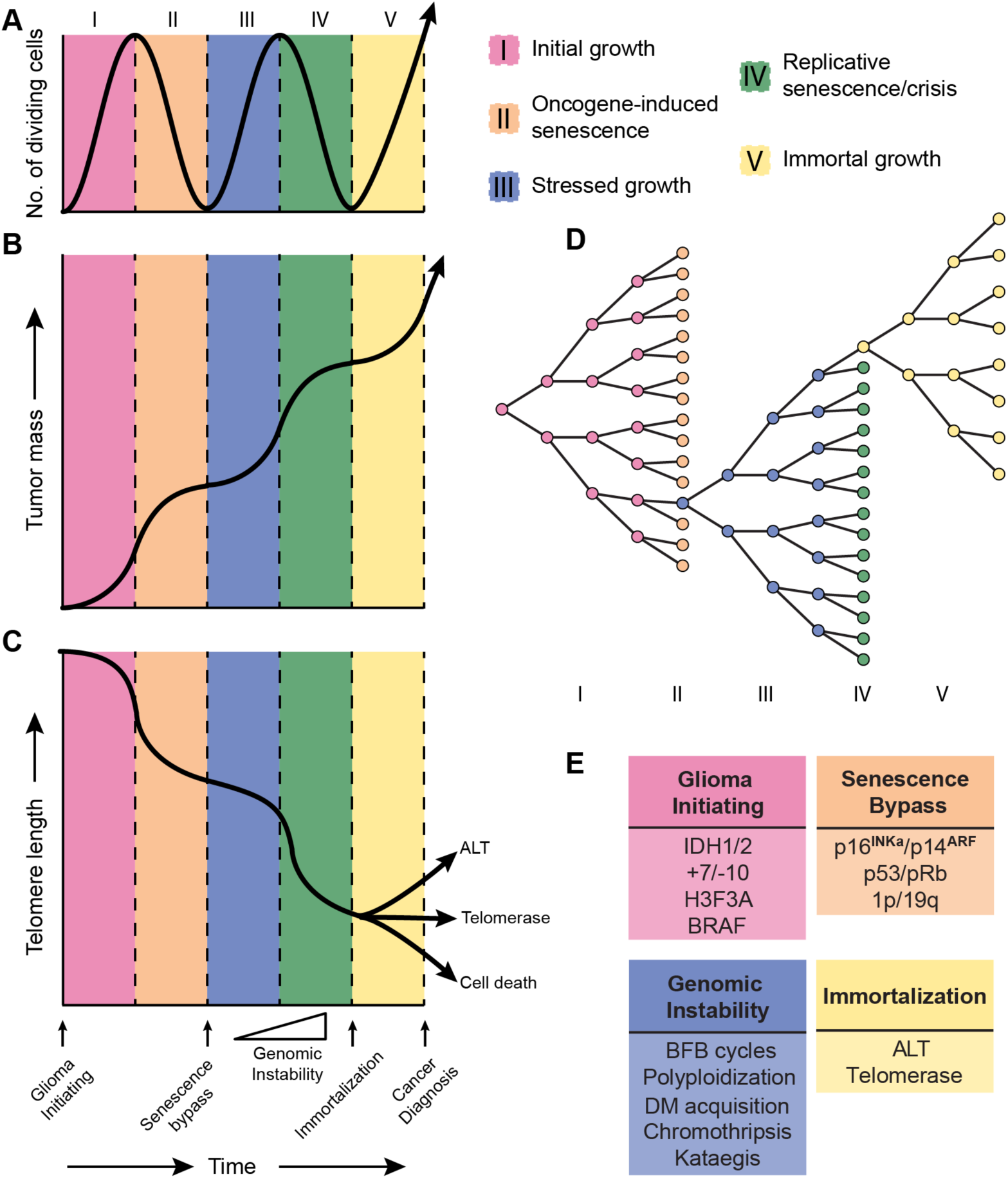
Model of the molecular life history of gliomas. The temporal sequence of events can be subdivided in five phases (I – V) represented in different colors. **A.** The number of dividing cells (or proliferation rate) across each phase. Proliferation peaks towards the end of growth phases and dips going into senescence phases. **B.** The tumor mass across each phase. Tumor mass increases exponentially during growth phases and logarithmically during senescence phases. **C.** Telomere length across each phase. Telomere length over time follows a pattern that is inverse to tumor mass. **D.** Cell doubling diagram indicating the growth barriers (senescence phases) and resulting selection bottleneck. **E.** Somatic alterations associated with different phases in gliomagenesis. The timing of each event is indicated on the x-axis of panel C. Genomic Instability events are accumulated during phase III-IV. Of note, this model is a simplified representation of true gliomagenesis. The x-axis is not drawn to scale, in part because the duration of the phases likely varies from cell to cell and between various tumor types. Furthermore, the position of the curves is arbitrary as cells in a tumor may not be in sync. BFB = breakage-fusion-bridge, DM = double minute, ALT = Alternative lengthening of telomeres

The initial growth phase (phase I) follows the acquisition of a glioma initiating oncogenic event and is characterized by aberrant proliferation of pre-tumor cells. Continued oncogenic exposure may impede tumor growth and trigger a durable form of cell cycle arrest termed oncogene induced senescence (phase II) in a majority of tumor cells. Some phase I/II cells may acquire molecular changes to bypass oncogene-induced senescence and continue growth in spite of unfavorable and stressful conditions including DNA damage and dysfunctional telomeres. Continued growth despite incremental genomic instability marks the third phase (phase III). This second round of glioma cell growth under harsh conditions triggers a second round of durable growth arrest termed replicative senescence (phase IV) and in some cases, brings forth a state of cellular crisis characterized by widespread cell death. Rare cells may acquire stem-like characteristics and a means to continue growth indefinitely, giving rise to a final phase of immortal tumor growth (phase V). Such phase V glioma stem-like cells uphold the tumor progenitor cell population *via* their capacity for self-renewal and may also give rise to more differentiated and growth arrested stage IV cells, losing their stem-like properties.

## 2. Classification of human gliomas

Gliomas encompass a very diverse group and account for the great majority of tumors originating in the parenchyma of the CNS [9]. Traditionally, gliomas are classified according to their microscopic similarities with (precursors of) glial cells and then designated as astrocytomas, oligodendrogliomas, mixed gliomas (mainly oligoastrocytomas) or ependymomas. Two larger glioma groups are recognized: so-called diffuse gliomas, characterized by extensive infiltrative growth into the surrounding CNS parenchyma, and more circumscribed (non-diffuse) gliomas such as pilocytic astrocytoma and ependymomas (Figure 1).

Diffuse gliomas are by far the most frequent gliomas in adult patients and histologically may show astrocytic, oligodendroglial or mixed phenotype of the tumor cells. After assessment of the glioma type, a malignancy grade is assigned to these tumors based on presence/absence of especially marked mitotic activity, florid microvascular proliferation (MVP), and necrosis. Diffuse gliomas in which these features are lacking are graded as low-grade (WHO grade II), tumors showing brisk mitotic activity in the absence of necrosis and florid MVP as anaplastic (WHO grade III) astrocytoma, oligodendroglioma, or oligoastrocytoma. The additional presence of necrosis and/or florid microvascular proliferation leads to a diagnosis of the most malignant astrocytic tumor, i.e. glioblastoma (previously designated as glioblastoma multiforme because of the wide range of histological features). In pure oligodendroglial tumors the presence of necrosis and florid MVP is not enough to consider the neoplasm as WHO grade IV [10].

For over a century, such microscopic evaluation has provided the gold standard for the diagnosis of gliomas, assessment of prognosis and formed the basis for therapeutic management. However, multiple studies showed that a purely histopathologic classification suffers from considerable inter- and intraobserver variability [11-13]. In the course of the last two decades it became increasingly clear that molecular characteristics may provide a more robust and objective basis for subtyping of diffuse gliomas. Since 2008, it has been demonstrated that in adult patients with diffuse gliomas based on the presence or absence of mutations in the isocitrate dehydrogenase 1 (*IDH1*) or *IDH2* gene and of complete, combined loss of the short arm of chromosome 1 and of the long arm of chromosome 19 (complete 1p/19q-codeletion) three major, clinically relevant subgroups can be defined, and both scientists and clinicians increasingly turned towards molecular markers to aid diagnosis [14-19].

Indeed, the International Society for Neuropathology – Haarlem Consensus Guidelines and the subsequently published, revised 4th edition of the WHO classification of CNS tumors (published in 2016) embrace the notion of an integrated histo-molecular classification of diffuse gliomas [20, 21]. The WHO classification nowadays recognizes the following three major molecular diffuse glioma subgroups:

- IDH-wildtype: most of these histologically represent astrocytic tumors, a large percentage belonging to the highest malignancy grade, i.e. glioblastomas
- IDH-mutant and 1p/19q-non-codeleted: these tumors also generally have an astrocytic phenotype, but a much larger percentage is at first diagnosis histologically lower grade/WHO grade II or III
- IDH-mutant and 1p/19-codeleted: most of these are characterized by a prominent oligodendroglial phenotype of the tumor cells

In this new classification, the presence of complete 1p/19q-codeletion in an IDH-mutant diffuse gioma is considered pathognomonic for oligodendroglioma, while IDH-mutant oligodendroglioma-appearing tumors lacking this event are reclassified as astrocytoma, IDH-mutant. Following this strategy, the diagnosis of mixed glioma/oligoastrocytoma can be expected to largely disappear, except when additional molecular tests cannot be performed or do not provide unequivocal results; in that situation, not-otherwise-specified (NOS) should be added to the diagnosis to indicate that ideally such samples require further workup. Rare IDH-mutant cases may show a dual genotype harboring both a 1p/19q-codeleted and non-codeleted component [22].

Diffuse midline glioma, H3 K27M-mutant, was added in the WHO 2016 classification as a separate entity as well [23-25]. These diffuse midline gliomas generally occur in children and, as the name implies, typically are located in the ’midline’ of the CNS (brainstem, thalamus, cerebellum and spinal cord), with diffuse intrinsic pontine glioma as a frequent representative. These H3-mutant diffuse midline gliomas are highly aggressive (WHO grade IV). By far the most frequent astrocytic tumors in children though are pilocytic astrocytomas. In sharp contrast to diffuse midline gliomas, pilocytic astrocytomas generally are more circumscribed (therefore grouped under non-diffuse gliomas) and show an indolent, WHO grade I behavior [26-28]. Meanwhile, the transition from a purely histological to a histo-molecular classification of especially diffuse gliomas represents a paradigm shift and necessitates re-evaluation of histologic criteria used for grading and guidance of therapeutic decisions [29, 30].

## 3. A Model for the Temporal Molecular Pathogenesis of Gliomas

### 3.1 Phase I: Initial Growth

The theory that cancer results from accumulation of mutations over time, in a subset of patients combined with contribution of inherited risk factors, has been around for over six decades and has been refined over the years [31-34]. For the purpose of this review we will consider a glioma initiating event to be the first acquired (somatic) event towards developing glioma. This event should provide a competitive growth advantage, either by directly increasing proliferation or by creating the conditions in which increased proliferation may happen. Cells and their progeny characterized by such an event will be primed to outcompete neighboring cells giving rise to an initial tumor mass. While germline events that contribute to glioma risk may precede such glioma initiating events, their incomplete penetrance suggests that they cannot be considered as causal for glioma formation and instead prime the environment for tumor formation [35]. Furthermore, even if a glioma initiating event marks the first somatic event in the formation of a tumor it may not be responsible for initiating growth directly. Instead, this event may promote tumorigenesis indirectly *via* stochastic activation of oncogenes or repression of tumor suppressors. Genetic or epigenetic selection pressures will prioritize daughter cells with growth advantages over those without and daughter cells with lethal genotypes will rapidly disappear [36-38].

#### 3.1.1 IDH-mutant diffuse gliomas

Mutations in *IDH1*/*IDH2* are commonly considered to be glioma initiating. Several studies have shown that they are amongst the few alterations highly shared amongst primary and recurrent tumors [39-41] which is explained by their presence in the cell of origin and all cells derived thereof. Comparing multiple biopsies from the same tumor, IDH mutation events fit the proposed criteria of a glioma initiating event [42-44]. Laboratory experiments have demonstrated that IDH mutations alone are sufficient to reprogram the transcriptome and epigenome of normal cells to prevent these cells from entering a terminally differentiated state [45-47].

IDH dysregulation likely contributes to gliomagenesis *via* the accumulation of the oncogenic metabolite R(-)-2-hydroxygulatarate (2HG) [48, 49]. Wildtype IDH enzymatically converts isocitrate into α-ketoglutarate (α-KG) as part of the citric acid cycle, whereas mutant IDH metabolizes α-KG into 2HG [50, 51]. Both mutant and wildtype IDH alleles are therefore essential for the oncogenic function of IDH. IDH mutations in glioma result in genome-wide hypermethylation [46, 52], most likely due to effects of 2HG on the Ten-eleven translocation methylcytosine dioxygenase (Tet) family of proteins [45, 53, 54]. This hypermethylation may provide a growth advantage to cancer cells due to the epigenetic activation of oncogenes *via* stochastic activation of alternative gene regulatory programs, some conferring added fitness [36]. One such mechanism in glioma may be *via* methylation induced disruption of a CCCTC-binding factor (*CTCF*) binding site, resulting in aberrant activation of Platelet-Derived Growth Factor Receptor Alpha (*PDGRFA*) [55].

#### 3.1.2 IDH-wildtype diffuse astrocytomas

Approximately 70% of IDH-wildtype diffuse astrocytomas are characterized at the molecular level by a single copy loss of chromosome 10 and gain of chromosome 7 (+7/-10) [17, 56]. Loss of chromosome 10, and of chromosome arm 10p in particular have been reported to occur more frequent and may precede gain of chromosome 7 and loss of 10q in mutational time [57]. Based on evolutionary modeling using primary-recurrent tumor pairs and multisector tumor sampling, several independent groups have found that +7/-10 is homogeneous and longitudinally preserved and thus likely the first and tumor-initiating event in a large fraction of IDH-wildtype diffuse astrocytomas/glioblastomas [44, 58-61]. A recent study suggested that gains of chromosome 7 likely occur in the first 10% of mutational time and are preceded by loss of chromosome 10, consistent with existing data [61]. In the past several years there has been a lot of interest in *TERT* promoter mutations and an increasing body of evidence suggests that these mutations precede +7/-10 [62]. Nevertheless, a potential role for *TERT* promoter mutations to promote proliferation in the initial growth phase and as a tumor-initiating event is speculative and will be discussed later in this review.

Chromosome 7 is home to several oncogenes that have been implicated in gliomagenesis such as Cyclin Dependent Kinase 6 (*CDK6*), MET Proto-Oncogene (*MET*) and Epidermal Growth Factor Receptor (*EGFR*) while chromosome 10 hosts several tumor suppressor genes, including Tet family member Tet Methylcytosine Dioxygenase 1 (*TET1*) and Phosphatase and Tensin Homolog *(PTEN)*. Likely these genes alone are not by themselves responsible for initiating glioma formation, and combined chromosome 7 gains and 10 losses initiate IDH-wildtype gliomagenesis via complex dosage-dependent modulation of a multitude of tumor promoting and suppressing genes [63].

Some IDH-wildtype diffuse gliomas show cytogenetically intact chromosomes 7 and 10, implying that other initiating events may give rise to these tumors. Such events may include activating or inactivating alterations in the phosphoinositide 3-kinases (PI3K), tyrosine kinase receptor (RTK) and mitogen-activated protein kinase (MAPK) pathways [64]. PI3K-pathway alterations include mutations in PI3-Kinase Subunit Alpha (*PIK3CA*), PI3-Kinase Subunit P85-Alpha (*PIK3R1*), or inactivation the aforementioned tumor suppressor *PTEN* [65-67]. Glioma-initiating events may also include point mutations in RTK-pathway genes such as *EGFR* and *PDGFRA* or in MAPK-pathway genes such as Neurofibromin 1 (*NF1*) [68]. Much is already known about the effect of these mutations on cancer growth but additional research is needed to secure their potential role as glioma initiating events.

A particular subgroup of diffuse gliomas is characterized by mutations in H3 histone family members and these gliomas occur most often in children [23, 69, 70]. The diffuse midline glioma, H3 K27M-mutant, shows a lysine to methionine substitution at position 27 of the *H3F3A* gene and is included in the WHO 2016 classification as a separate entity. Other H3-mutant diffuse gliomas in children and adolescents occur predominantly in the cerebral hemispheres and often show H3 G34R/V mutation (implying a glycine 34 to arginine or valine substitution) [69-72]. In contrast to hypermethylated IDH-mutant gliomas, these H3-mutant gliomas show a general DNA hypomethylation phenotype [73]. A recent study used *in vivo* and *in vitro* models to show that tumors with a K27M variant demonstrated significantly lower rates of histone 3 with tri-methylated lysine 27 (H3K27me3) and the mutation inhibited the Polycomb repressive complex 2, a protein complex associated with long term epigenetic silencing of developmental genes [74]. Taken together these findings suggest that H3 K27M and G34R/V variants are glioma initiating events that contribute to tumorigenesis *via* stochastic activation of developmental genes and an epigenetic selection process similar to what has been proposed in the context of IDH mutations [36, 75].

#### 3.1.3 Non-diffuse gliomas

Recent studies have shown that pilocytic astrocytomas near universally harbor abnormalities in the MAPK-pathway, and most commonly a tandem duplication targeting chromosome 7q, which gives rise to a *KIAA1549-BRAF* fusion gene consisting of the N-terminus of *KIAA1549* and the kinase domain of v-RAF murine sarcoma viral oncogene homolog B1 (*BRAF*) [28]. Alternative alterations include the oncogenic V600E missense mutation also targeting *BRAF* [26]. The *BRAF* V600E mutation results in an activating change due to a substitution of valine with glutamic acid at codon 600. In a non-cancer setting, *BRAF* activates kinases MEK and ERK, which in turn activate transcriptional machinery to promote differentiation, proliferation, growth and apoptosis [76]. Both *BRAF* V600E mutations and *BRAF* fusion genes contribute to tumorigenesis by constitutively activating the kinase domain of *BRAF*, resulting in overactive signaling activity and a selective growth advantage for affected cells [77-79]. In most pilocytic astrocytoma (even after thorough analysis) an activating change in *BRAF* or other MAPK-pathway members is the only genomic change that can be confidently detected, implying that it is the glioma initiating event in this disease [80].

### 3.2 Phase II: Oncogene-induced senescence

Continued oncogenic signaling in the initial growth phase prompts the activation of tumor suppressive signaling *via* activation of the p16^INK^4^a^/p14^ARF^-RB-p53 cell cycle and cell stress pathways (Figure 3), slowing tumor growth and transitioning a majority of cells with intact pathways into a terminal state called oncogene-induced senescence [81]. First discovered over five decades ago in cultured fibroblasts, senescence is a stress-induced durable cell-cycle arrest [82]. The role of senescence in cancer has been reviewed extensively [83-88]. Briefly, senescence provides a major tumor suppressive barrier and dividing tumor cells are put under selection pressure to acquire molecular events to prevent its onset. Hallmarks of senescence include durable growth arrest; short, dysfunctional telomeres; and a marked increase in DNA damage and stress signaling [85]. Although senescent cells are growth arrested, they are metabolically active and release a plethora of signaling molecules to the microenvironment, also known as the senescence-associated secretory phenotype [88]. Distinction must be made between oncogene-induced senescence which is discussed here and is triggered by chronic oncogenic signaling, and replicative senescence discussed later, describing senescence triggered by telomere dysfunction following extensive replicative cycles [86]. Oncogene-induced senescence poses a significant growth barrier, and most cells will not acquire molecular alterations that allow them to bypass this barrier and will therefore become senescent [81]. However, rare cells may acquire such alterations, eliciting a selective sweep by a subclone that will rapidly dominate the tumor cell population.

**Figure 2.**
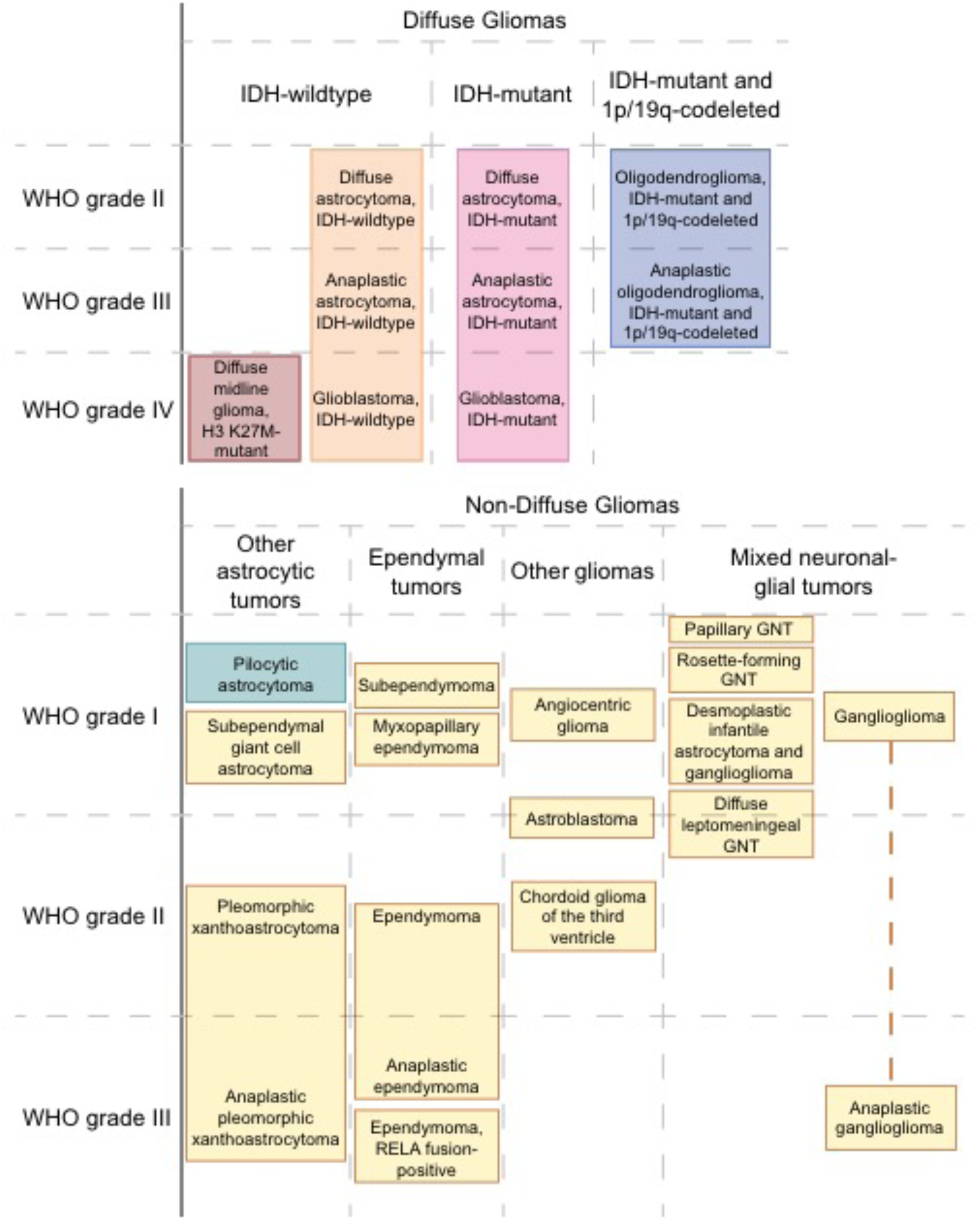
Schematic of the 2016 WHO classification of diffuse and non-diffuse gliomas. Note that especially diffuse gliomas are nowadays classified based on integration of histologic and molecular characteristics. This review focusses on diffuse gliomas, and on pilocytic astrocytomas (the remaining glioma categories are shaded in yellow). Diffuse gliomas are graded as WHO grade II to IV (least to most malignant). Many non-diffuse gliomas are relatively indolent (WHO grade I), while the most malignant tumors in this category are graded as WHO grade III. For some rare and/or only recently identified neoplasms (esp. angiocentric glioma, diffuse leptomeningeal glioneuronal tumor) an unequivocal malignancy grade has not been assigned yet.

**Figure 3.**
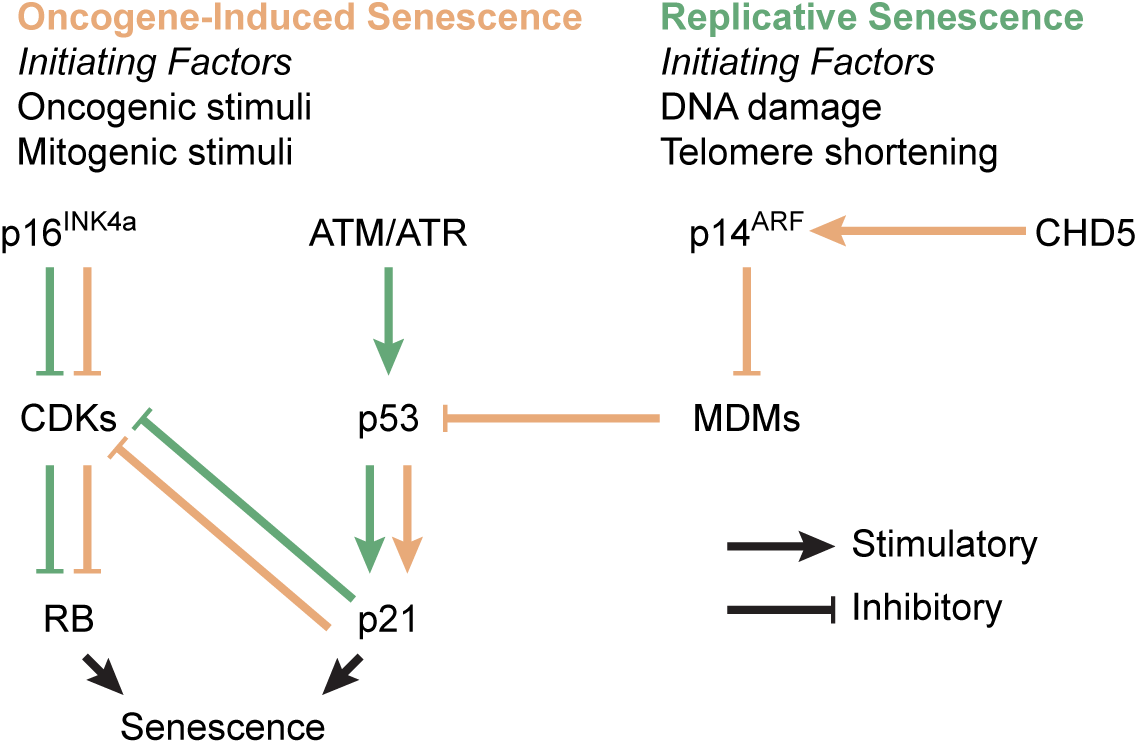
Process diagram indicating the p16^INK4a^/p14^ARF^-RB-p53-pathway in normal conditions. Disruption of one or multiple components through mutation or copy number change may prevent or suppress the onset of senescence. Various stimuli use different routes to activate the senescence response, leaving compensatory mechanisms in place in case components fail. For example, if oncogene-induced senescence is repressed via *CDKN2A/B* inactivation, DNA damage and telomere shortening could still trigger replicative senescence via *ATM* and *ATR*. CDKs = Cyclin Dependent Kinases (eg. CDK2), MDMs = Murine Double Minutes (eg. MDM2)

In some cancer types a senescent precursor stadium can be identified, such as intestinal polyps in colon cancer and dysplastic naevi in melanoma [89]. In contrast to the catastrophic karyotypes demonstrated by their later stage derivatives, these growth-arrested senescent precursor lesions entered a senescent state *via* the activation of single oncogene such as *BRAF* and are otherwise genetically unremarkable [90-93]. This begs the question if such premalignant precursor lesions exist for diffuse gliomas as well. If so, they may however well be microscopic in size and go unnoticed in imaging or autopsie studies.

The tumor suppressor proteins p16^INK^4^a^, p14^ARF^, RB and p53 can be considered as gatekeepers of senescence. The INK4a/ARF locus on chromosome 9 contains both Cyclin Dependent Kinase Inhibitor 2A (*CDKN2A*) and 2B (*CDKN2B*), combined encoding for both p16^INK^4^a^ and p14^ARF^ *via* alternative splicing. Tumor suppressors RB and p53 on the other hand are encoded for by the genes Retinoblastoma 1 (*RB1*) and Tumor Protein 53 (*TP53*) on chromosomes 13 and 17, respectively. Inactivation of one or multiple of these genes *via* genomic deletion and/or inactivating mutations has been linked to repression of senescence signaling and are a common event in all cancers including gliomas [83, 94-96]. For the purpose of this review, we define the term ‘senescence bypass event’ as any molecular alteration that suppresses the onset of oncogene-induced senescence.

#### 3.2.1 IDH-mutant diffuse astrocytomas

IDH-mutant astrocytomas are often characterized by loss of one allele of *TP53*, combined with a loss-of-function mutation in the remaining allele. The frequency of *TP53* mutations in IDH-mutant (’secondary’) glioblastomas is comparable to that in lower grade IDH-mutant astrocytomas from which these glioblastomas are derived via malignant progression, suggesting that *TP53* aberrations are early lesions in these tumors [97]. Furthermore, and in contrast to IDH-wildtype gliomas, *TP53* mutations were shared between all *TP53*-mutant cases of primary and recurrent tumors in a recent study [39]. Analysis of multiple biopsies from the same IDH-mutant tumors indicated that samples mutant for *TP53* were always IDH mutant, while some IDH-mutant samples lacked *TP53* mutations, suggesting that IDH mutations precede *TP53* inactivation [98]. IDH mutation and *TP53* inactivation therefore both comprise early events in gliomagenesis, with *TP53* inactivation generally following mutation in IDH. The p53 tumor suppressor protein is involved in many different functions, and especially its role in cell cycle arrest and senescence is very well understood [99]. While enzymatically active wildtype p53 triggers senescence in response to oncogenic stress, mutant p53 inadequately blocks proliferation thereby preventing the onset of oncogene-induced senescence. In addition, loss of p53 enzymatic activity circumvents replicative senescence and triggers crisis, as discussed in phase IV [94].

#### 3.2.2 IDH-mutant oligodendrogliomas, 1p/19q-codeleted

The majority of IDH-mutant tumors wildtype for *TP53* demonstrate a combined single copy loss of the complete chromosome arms 1p and 19q (complete 1p/19q-codeletion) [5, 100]. These tumors are according to the revised WHO criteria the canonical oligodendrogliomas [21]. 1p/19q-codeletions were found to be stable across longitudinal samples and multiple biopsies, suggesting that they are early events [101-103]. The finding that codeleted tumors are almost exclusively IDH-mutant, while the reverse is not true, suggests that mutations in IDH precede codeletion. The origin and mechanism behind 1p/19q-codeletions remain to be resolved. Additional loss-of-function mutations in far-upstream element binding protein (*FUBP1*) on 1p31.1 and capicua transcriptional repressor (*CIC*) on 19q13.2 are observed in over 60% of 1p/19q-codeleted gliomas and may serve as second hits although their functional significance remains unclear [104]. A minor adverse survival association was found for *FUBP1* mutant tumors in one study, supporting a role for these mutations in tumorigenesis [105]. Codeletion of 1p and 19q may allow tumors to bypass oncogene-induced senescence through the mono-allelic inactivation of tumor suppressor genes on these chromosome arms [63]. One mechanism to achieve this may be through a dosage-dependent repression of the gene Chromodomain Helicase DNA Binding Domain 5 (*CHD5*) on 1p36 [106-108]. A study that used genetic engineering to create mouse models with gains and losses of a region corresponding to human 1p36 housing *CHD5* found that duplication of 1p36 led to decreased proliferation and senescence whereas a single-copy deletion led to immortalization [108]. This study further shows that its protein product Chd5 regulates p16^INK^4^a^ and that loss of Chd5 compromises p53 function, providing direct evidence that this gene is involved in senescence regulation. Another gene target may be the p53 homolog Tumor Protein 73 (*TP73*), which also resides on 1p36, directly interacts with p53 and may activate p53 target genes [109]. Most likely, as may be the case with growth-promoting +7/-10 events observed in IDH-wildtype gliomas, 1p/19q-codeletions may act on senescence bypassing pathways by modulating a multitude of tumor suppressor genes.

#### 3.2.3 IDH-wildtype diffuse astrocytomas

Amongst IDH-wildtype astrocytomas/glioblastomas, one of the most frequent alterations is a homozygous loss of *CDKN2A* and *CDKN2B* [110]. Mathematical modelling has suggested that homozygous *CDKN2A/B* loss occurs after +7/-10 but before other molecular events [59]. Homozygous *CDKN2A/B* loss alone is insufficient for tumor formation in mice, requiring the activation of an oncogene to generate tumors *in vivo* [111]. As such, homozygous *CDKN2A/B* loss is likely a second event in the tumorigenesis of IDH-wildtype astrocytoma or glioblastoma. The role of protein products p16^INK^4^a^ and p14^ARF^ in senescence are very well understood. Indeed, *CDKN2A/B*^NULL^ astrocytes can grow indefinitely in culture [112], and introduction of p16^INK^4^a^ in immortal human glioma cell lines leads to cell cycle arrest and senescence [113]. Mutations in *TP53* sometimes co-occur with homozygous *CDKN2A/B* loss in IDH-wildtype glioma [114]. Where *TP53* mutations can be found across all tumor cells in IDH-mutant astrocytoma, *TP53* mutations can often only be detected in a subset of tumor cells amongst IDH-wildtype astrocytoma [44]. Furthermore, *TP53* can be counted as the top gene whose status changes in recurrent compared to primary IDH-wildtype glioblastoma. Both a loss of and a gain of a mutant *TP53* allele have frequently been described in recurrent IDH-wildtype tumors. These observations indicate some functional redundancy between alterations in p53 and p16/p14 in IDH-wildtype glioma. Pediatric H3-mutant/IDH-wildtype diffuse gliomas are for the most part *TP53* mutant while *CDKN2A/B* remain intact [23].

#### 3.2.4 Non-diffuse gliomas

Several lines of evidence suggest that pilocytic astrocytomas (WHO grade I, IDH-wildtype) are arrested in a senescent phase II state and do not advance to later phases. First, these tumors were found to frequently demonstrate several biomarkers of senescence at tumor detection, including widespread β-galactosidase activity and p16^INK^4^a^ staining [115-117]. Second, these tumors are very quiescent genetically, often demonstrating but a single activated oncogene, such as a *BRAF* fusion, *BRAF* V600E mutation or rarely an activating mutation in *FGFR1* or *PTPN11* [118-120]. Third, these tumors grow slowly, harbor excellent outcomes and sometimes regress, perhaps because these tumors do not immortalize [121-124]. Fourth, expression of activated *BRAF* V600E alone does not lead to tumor development in *in vitro* and in *in vivo* mouse models, while the combined activation of *BRAF* and loss of *CDKN2A/B* is transforming, suggesting that additional mutations to bypass senescence are required to advance to phase III [125-127].

### 3.3 Phase III: Stressed Growth

Cells presenting with continued proliferative signaling beyond the oncogene-induced senescence barrier are generally characterized by defective DNA damage response signaling and continued growth in a stressed environment. During this phase the repetitive DNA at the telomeric terminal ends of chromosomes become increasingly important [128]. Telomeres become progressively shorter as cells divide due to the linear conformation of chromosomes and directional replication machinery, a phenomenon that is critically important for diseases like cancer which are characterized by often rampant proliferation [129]. Telomeric DNA takes on a lasso conformation called the t-loop, and these loops are bound by the shelterin DNA binding protein complex. Together, these characteristics protect chromosome ends from being recognized as DNA double strand breaks and prevent inadvertent activation of DNA damage response pathways [130].

Dysfunctional telomeres are critically short and improperly protected telomeres lacking t-loops and shelterin complexes. They trigger the activation of DNA damage response pathways via ataxia telangiectasia mutated (*ATM*) and ataxia telangiectasia and Rad3-related (*ATR*) kinase, which are triggered by exposed and unprotected double stranded and single stranded DNA break ends, respectively. The exposed ends then fall victim to homology directed repair (HDR) and non-homologous end joining (NHEJ) repair processes, intended to repair accidental DNA breaks but lead to gross genomic instability when triggered by dysfunctional telomeres. When telomeres are unprotected, these repair processes prompt sister chromatids to fuse with one another, forming a dicentric chromosome. During the anaphase, the dicentric chromosome will form a bridge spanning the mitotic spindle and connecting the two daughter cells. Resolution of the chromatin bridge via cytoplasmic 3’ nuclease *TREX1* results in breakage of the dicentric chromosome at a locus not necessarily at the site where the fusion had occurred, resulting in an unbalanced inheritance of genetic material between the two daughter cells. Because the resulting daughter cells also lack telomeres, this process of breakage-fusion-bridge (BFB) cycles (Figure 4A) will repeat itself every subsequent cell division until telomeres are restored [131] [132-135]. The detrimental genomic instability acquired via telomere dysfunction and BFB-cycles endows these cells with powerful stochastic mutator mechanisms to acquire changes that provide a survival benefit under selective pressure.

**Figure 4.**
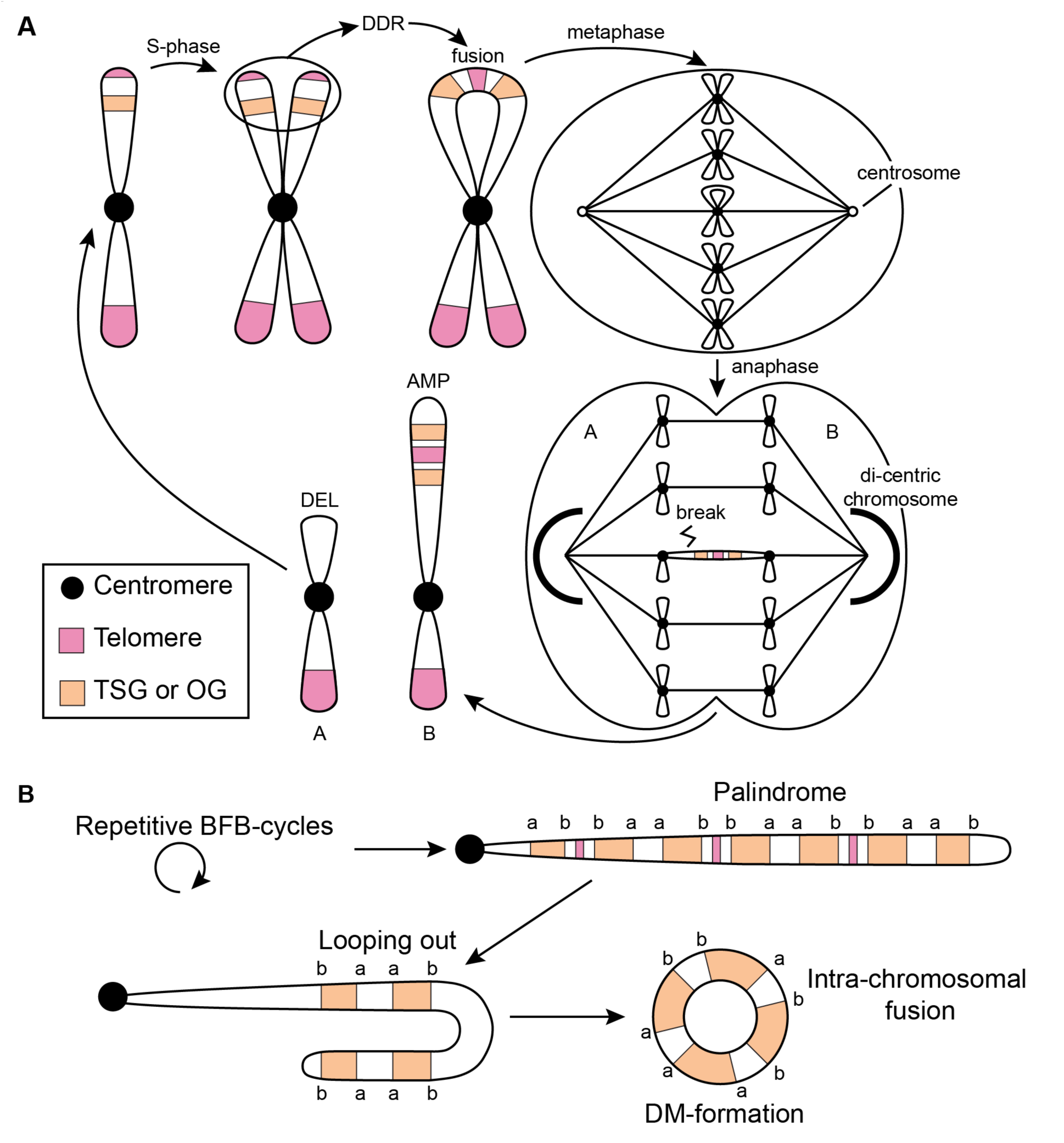
Genomic instability related to telomere stress **A.** Schematic illustrating BFB-cycles. Following a single BFB-cycle, daughter cells are left with unequal DNA content, leading to a deletion in A and an amplification in B. BFB cycles may also involve fusion of non-sister telomeres (not shown). **B.** Repetitive BFB-cycles form palindromes demonstrating high intra-segmental homology. This can lead to intra-chromosomal fusions and formation of double minutes. TSG = tumor suppressor genes; OG = oncogene; DDR = DNA damage response; AMP = amplification; DEL= deletion; DM = double minutes; BFB = breakage-fusion-bridge

The intensity of genomic instability endured during the stressed growth phase may depend on the severity of senescence pathway dysregulation incurred overcoming oncogene-induced senescence (Figure 2). In the case of H3-mutant, IDH-wildtype and IDH-mutant astrocytoma/glioblastoma this pathway is perturbed close to the source via the direct loss of RB, p53 or p16^INK^4^a^ protein function. In IDH-mutant and 1p/19q-codeleted oligodendroglioma this pathway may be repressed indirectly, for example *via* the modulation of p14^ARF^ activity through a partial deletion of *CHD5*. This may explain why the latter group of tumors show significantly less genomic instability compared to the former tumor types. Moreover, in pilocytic astrocytoma this pathway may not be affected at all and these tumors may not advance beyond oncogene-induced senescence. While genomic instability incurred during this phase demonstrates some similarities within glioma subtypes, there appears to be a striking degree of entropy shared between glioma subtypes and we therefore did not separate this section according to tumor type. More research is needed to carefully delineate the selective pressures at play to better understand differences and similarities between various glioma entities.

The advent of high-throughput sequencing has led to remarkable progress in understanding the complexity of genomic instability in cancer, including complex deletions, amplifications and translocations [38]. It is important to note that the genomic organization that can be reconstructed using sequencing at the time of analysis are those changes that resulted in viable cells and were selected for. Recent work has shown that telomere dysfunction directly leads to catastrophic genomic events, including genome shattering (chromothripsis), clustered regions of focal hypermutation (kataegis) and whole genome doubling (tetraploidization) [136-138]. Polyploidization may occur both before or after chromothripsis, suggesting that these may occur in a similar stage of tumor development [139]. Comparison of primary and relapsed tumors across various tumor types including gliomas demonstrated that relapsed tumors lack additional catastrophic events, suggesting that these occurred during a stressful period the initial development of the tumor that was later stabilized [137].

Recently there has been a renewed interest in circular extrachromosomal DNA elements called double-minute (DM) chromosomes in cancer, and it was shown that such DMs are frequent in glioma [140, 141]. Although DM chromosomes have been long recognized as a cytogenetic feature of cancer, relatively little was known about its biological relevance. DM chromosomes have a predisposition to involve cancer oncogenes such as MYC Proto-Oncogene Protein (*MYC)*, MDM2 Proto-Oncogene (*MDM2*) or Cyclin Dependent Kinase 4 (*CDK4)* [142-146]. A unique feature of DM is that they lack centromeres and telomeres to dictate the organization of the mitotic spindle during mitosis, and are therefore randomly distributed across daughter cells [147]. Interestingly, this feature hypothetically provides DM chromosomes with an impressive fitness advantage over linear chromosomes as they do not need telomeres to protect them from inadvertent DNA damage response pathways and are not subjected to detrimental BFB-cycles. It has been proposed that DM chromosomes are the result of the fusion and circular assembly of linear DNA segments arising from BFB-cycles (Figure 4B) [148, 149]. These findings suggest that DM-chromosomes may be positively selected for during telomere dysfunction. Compared to IDH-mutant gliomas, DMs in IDH-wildtype tumors more often involve established glioma oncogenes, despite what appears to be a comparable frequency of DMs in both glioma categories [140]. More research is needed to precisely determine the frequency of DMs in glioma subtypes and to pinpoint the genetic origin of these structures.

It was recently shown that upon recurrence of IDH-mutant tumors the IDH locus may be deleted in some tumors [150]. Furthermore, IDH-mutant tumors are very hard to culture, and when it succeeds, IDH mutations that were present initially have been reported missing, raising the possibility that losing an IDH mutation is advantageous for survival in culture [151]. Introduction of mutant IDH in cell cycle checkpoint deficient cells rapidly transforms these cells into competent tumor cells [152]. However, IDH inhibition in these cells after as little as four days after its first introduction did little to slow tumor growth. These findings suggest that IDH-mutant tumors rapidly evolve and acquire new driver events that uphold the tumor sustaining population and make the initial IDH mutation redundant. Similarly, complex structural alterations affecting known glioma drivers such as *EGFR, PDGFRA* and MET Proto-Oncogene, Receptor Tyrosine Kinase (*MET*) were found to be late events not shared across primary/recurrent tumor pairs or multiple sampling from single tumors [44, 60].

Chromothripsis is rare, DM-chromosomes and poliploidization are more common and BFB-cycles are likely nearly universally present in diffuse gliomas. The etiology of these mutator mechanism is for the most part unknown and more research is needed to understand why the frequency of the events varies between glioma subtypes. Telomere dysfunction and stressed growth may thus promote the context-dependent evolution of glioma cells, rendering glioma initiating events redundant and providing gliomas with new fuel that rapidly increase intratumoral heterogeneity and can deal with various toxic stresses and bottlenecks. While stochastic mutator mechanisms in the stressed growth phase provide ample selection pressure to acquire beneficial changes, the detrimental genomic instability under which cells must operate acts as a powerful tumor suppressive barrier. Unchecked growth will rapidly lead to another round of DNA-damage induced replicative senescence, or when those checkpoints fail completely, cell crisis.

### 3.4 Phase IV: Replicative senescence/crisis

Sustained stressed growth is not durable and will eventually lead tumor cells down to one of two possible roads. Tumor cells with a partially intact senescence response (i.e. functional p53 and RB) may undergo a second round of senescence called replicative senescence in response to dysfunctional telomeres. Tumor cells with a completely dysfunctional senescence response (i.e. loss-of-function mutation in *TP53* or *RB1*) instead continue proliferating in a state of cellular crisis leading to cell death in a vast majority of cells [153]. Replicative senescence and crisis both pose a second population bottleneck to further tumor formation. It is essential that tumor cells transition to a less stressful environment with proper telomere maintenance to prevent further BFB-cycles and other catastrophic events. Cell culture experiments have demonstrated that direct immortalization of cells prior to a stressed growth phase enables them to bypass genomic instability and immortalize lacking the wild karyotypes typically associated with malignant transformation [154, 155]. These observations suggest that genomic instability in cancer development precedes immortal growth and is required to generate errors enabling telomere maintenance [154].

Acquisition of a telomere maintenance mechanism endows cancer cells with immortal growth, meaning that they are bestowed a limitless replicative potential [156]. Telomere maintenance is established once a tumor cell has reactivated telomerase or activated alternative lengthening of telomeres. Moreover, restoration of telomere function may prevent further BFB-cycles and restore genome stability. The canonical pathway involves the reactivation of the ribonucleoprotein telomerase which is transcriptionally silent in differentiated adult cells [157]. The telomerase catalytic component telomerase reverse transcriptase (*TERT*) is expressed in over 80% of human cancers and is thought to be rate limiting for telomerase activity [158]. In the alternative pathway tumors become immortalized via a recombination driven mechanism called alternative lengthening of telomeres (ALT) [159].

#### 3.4.1 IDH-mutant diffuse astrocytomas

IDH-mutant diffuse astrocytomas almost universally demonstrate ALT [160]. ALT cells present with several defining characteristics, including a heterogeneous distribution of telomere length across chromosomes, extrachromosomal telomeric DNA fragments in a circular configuration (c-circles), increased expression of telomeric repeat-containing RNA (*TERRA*) from telomeres, the formation of ALT-associated promyelocytic leukaemia bodies (APBs), frequent telomere sister chromatid exchanges (T-SCEs) and recombination between telomeres from different chromosomes [161]. ALT provides cancer cells with stabilizing telomere maintenance in a telomerase negative setting. Although it remains unclear precisely how ALT becomes activated, its presence has been tightly associated to loss-of-function events targeting the a-thalassemia/mental retardation syndrome X-linked (*ATRX*) or death-domain associated protein (*DAXX*) genes and these events are also a hallmark feature of IDH-mutant astrocytomas [162-164]. *ATRX* functions as an ATP-dependent helicase within the SWI/SNF family and combined these two genes form the ATRX-DAXX complex, which functions as a histone chaperone to deposit the histone variant H3.3 at telomeres [165]. Telomeric DNA has a tendency to form secondary quadruplex structures that challenge the replication machinery and need to be resolved for proper replication [166]. Although it remains unclear how these two genes protect telomeres from recombination and ALT, it has been suggested that the combined helicase activity of *ATRX* and the histone chaperone capabilities of the ATRX-DAXX complex can resolve the secondary quadruplex structure at telomeres, thereby enabling proper progression of the replication fork during S-phase and preventing the inadvertent activation of recombination (ALT) mechanisms [167, 168].

The presence or absence of inactivating *ATRX* and *DAXX* mutations present a strong correlation with ALT in many tumor types including gliomas [23, 162, 169]. Recent *in vitro* studies have shown that knockout of *ATRX* alone is insufficient to cause ALT, however, *ATRX* knockout combined with inactivation of p53 and RB enzymatic activity led to an increased incidence of ALT after enduring several cycles of telomere induced crisis [170-172]. Furthermore, the reintroduction of *ATRX* expression in *ATRX* mutant ALT cells led to a repression of T-SCE, APBs and c-circle formation [170, 171].

#### 3.4.2 IDH-mutant oligodendrogliomas, 1p/19q-codeleted

Oligodendrogliomas, IDH-mutant and 1p/19q-codeleted almost universally use telomerase to maintain telomeres and virtually all of these tumors carry an activating *TERT* promoter mutation [62]. These *TERT* promoter mutations are amongst the most common non-coding mutations in cancer [62, 173-175]. Recurrent hotspot point mutations substitute a cytosine at -228 or -250 relative to the promoter to a thymine (C228>T or C250>T) to create a de novo e-twentysix (*ETS*) transcription factor binding site that recruits the ETS family member GA-binding protein alpha chain (*GABPA*) to activate transcription [176].

Although the timing of *TERT* promoter mutations is still under debate, several lines of evidence suggest that *TERT* promoter mutations arise early (phase I) in gliomagenesis, or perhaps even occur prior to the glioma initiating event. *TERT* promoter mutations preferentially occur in tissues with a lower rate of self-renewal and there are numerous reports on the extra-telomeric functions of *TERT*, including effects on the NF-κB and WNT/β-catenin pathway promoting tumor growth and invasiveness [62, 177, 178]. Combined this raises the possibility that these mutations may contribute to tumorigenesis via other pathways than its effect on telomerase alone, providing a biological reason for these mutations to contribute to gliomagenesis early in phase I and before the onset of dysfunctional telomeres. In a glioma-specific analysis it was found that nearly all tumors with the ‘phase I event’ +7/-10 or ‘phase II event’ 1p/19q-codeletion have *TERT* promoter mutations, whereas not all *TERT* promoter mutant gliomas have +7/-10 or 1p/19q-codeletions, which may indicate that *TERT* promoter mutations even precede +7/-10 and 1p/19q-codeletions [17]. Another group studied the mutation fraction and used multi-sector sequencing in 1p/19q-codeleted oligodendrogliomas and found that *TERT* promoter mutations indeed are early events and may occur before IDH mutations [42]. The idea that *TERT* promoter mutation occurs early is further corroborated by the finding that genetically engineered *TERT* promoter mutations in telomerase positive embryonic stem cells do not affect telomerase activity, however, upon differentiation these engineered cells remain telomerase positive and acquire immortality [179]. More recently, it was found that *TERT* promoter mutations in melanoma do not initially support telomere maintenance and telomeres shorten to critically short length despite harboring promoter mutations [180]. The effect on telomere length was not observed until later, which the authors attributed to a two-step immortalization process. One study even reported canonical (-228 and -250) somatic *TERT* promoter mutations in the blood of multiple non-cancer patients, indicating that these events could even occur before the onset of cancer and act to prime the tumor bed [181]. Although roles for *TERT* outside of telomere maintenance remain to be definitively understood, these observations provide a sound argument that *TERT* promoter mutations can occur early in or even before gliomagenesis while providing a means towards immortalization at a later stage.

#### 3.4.3 IDH-wildtype diffuse astrocytomas

The majority of IDH-wildtype diffuse gliomas use telomerase for telomere maintenance [182]. Re-analysis of previously published samples reclassified according to WHO 2016 criteria demonstrated that approximately 75% of diffuse IDH-wildtype gliomas, are *TERT* promoter mutant [183]. Thus, *TERT* promoter mutations are common in both the most (IDH-wildtype diffuse astrocytomas) and the least (IDH-mutant and 1p19q-codeleted oligodendrogliomas) aggressive diffuse gliomas, suggesting that *TERT* promoter mutations are not dictating their biological behavior. *TERT* promoter mutations may serve tumorigenesis slightly different in these two glioma subtypes, although it remains unclear how. It was further found that *ATRX* mutations occur in approximately 5% of IDH-wildtype diffuse astrocytomas [183]. The prevalence of ALT in IDH-wildtype diffuse gliomas is higher than the frequency of *ATRX* mutations, suggesting that some of these tumors may use ATRX-independent mechanisms to activate ALT [160]. In a similar fashion, the prevalence of telomerase activity is higher than the prevalence of *TERT* promoter mutations in this tumor type, suggesting that these tumors may use *TERT* promoter independent mechanisms for the reactivation of telomerase [182]. Several candidate mechanisms have been previously described in glioma, including *TERT* promoter methylation or *TERT* amplifications [184]. Contrary to adult IDH-wildtype diffuse glioma, H3-mutant malignant pediatric glioma frequently demonstrate ALT [185], and several studies reported frequent co-occurrence of *ATRX* mutations in both H3 K27 and G34 mutant gliomas. However, reports of co-occurrence vary between 30% and 60% for K27 and 75% to 100% for G34, suggesting that there is a role for telomerase in many of these tumors as well [186].

### 3.5 Phase V: Immortalization & Dedifferentiation

The glioma stem cell theory states that amongst all cancerous cells in a tumor, a subset of cells act as progenitor or stem cells with reproductive capabilities and sustaining the cancer, much like normal bone marrow stem cells are responsible for replenishing the population of circulating leukocytes [187]. It has often been contrasted to the theory of clonal evolution, which suggests that cancers evolve through an iterative process of clonal expansion of from a single cell [188]. Recent advances in single-cell sequencing and lineage tracing have unveiled multiple populations of tumor cells in bulk tumor samples, providing fuel for the cancer stem cell hypothesis [189-192]. One study used single cell RNA sequencing on IDH-mutant and 1p/19q-codeleted oligodendroglioma patient samples and uncovered distinct cell populations of undifferentiated tumor stem cells and cells that are more differentiated along various glial lineages [190]. In a similar study of IDH-mutant astrocytoma the authors were able to detect the same cellular populations but with a higher ratio of stem-like to differentiated cells that increased with increasing WHO grades [191]. Another study used DNA barcoding of repeatedly *in vivo* transplanted glioblastoma cells to trace the lineage during their engraftment and found a population of progenitor cells that sustained the tumor and gave rise to differentiated non-proliferative cells, confirming results on a cellular hierarchy with fewer stem cells and a higher number of cycling progenitor cells from single-cell RNA sequencing studies and demonstrating that a relatively large fraction of cells can be tumor-initiating [192].

These studies all provide support for a cancer stem cell hypothesis and raise the question how these findings fit with our model, which leans towards a model of clonal evolution. Current evidence suggests that both mechanisms are acting together (Figure 5A). Whereas clonal evolution is important to establish the initial cancer stem cell population, neutral evolution (in line with the cancer-stem cell hypothesis) may fit better once the initial cancer core has been established, especially so when further evolutionary stimuli (e.g. senescence barriers, hypoxia, treatment) are lacking. The concept of neutral evolution holds that most molecular changes are not caused by natural selection but rather by the stochastic allelic variation that are neutral and do not affect cellular fitness [193]. A recent study analyzed cancer genomes from TCGA and found neutral evolution in approximately one-third of a wide spectrum of over 900 tumors, including 35 gliomas of which 8 (23%) suggested evidence in support of a neutral evolution process [194]. The authors conclude that all clonal selection must have occurred before the onset of cancer growth and not in later arising subclones. Several groups have since challenged these findings and suggested that their analysis does not univocally prove neutral evolution starting from the first malignant cell [195-197]. While it may be possible that tumors evolve neutrally beyond the most recent selective sweep we do not agree with the conclusion that these findings suggest that all clonal selection must occur before the onset of tumor growth. The authors do not take into account that at the time the tumor presents itself and is surgically removed, all remnants of a selection process that have been outcompeted or died will have completely disappeared, in contrast to “neutral” variants which do not affect fitness and will remain. According to our simplified model, tumorigenesis sequentially follows phase I to V. Once the cancer stem cell population has been established, tumor cells will follow neutral evolution, as long as new evolutionary or other stimuli for opportunistic growth are lacking (Figure 5A).

**Figure 5.**
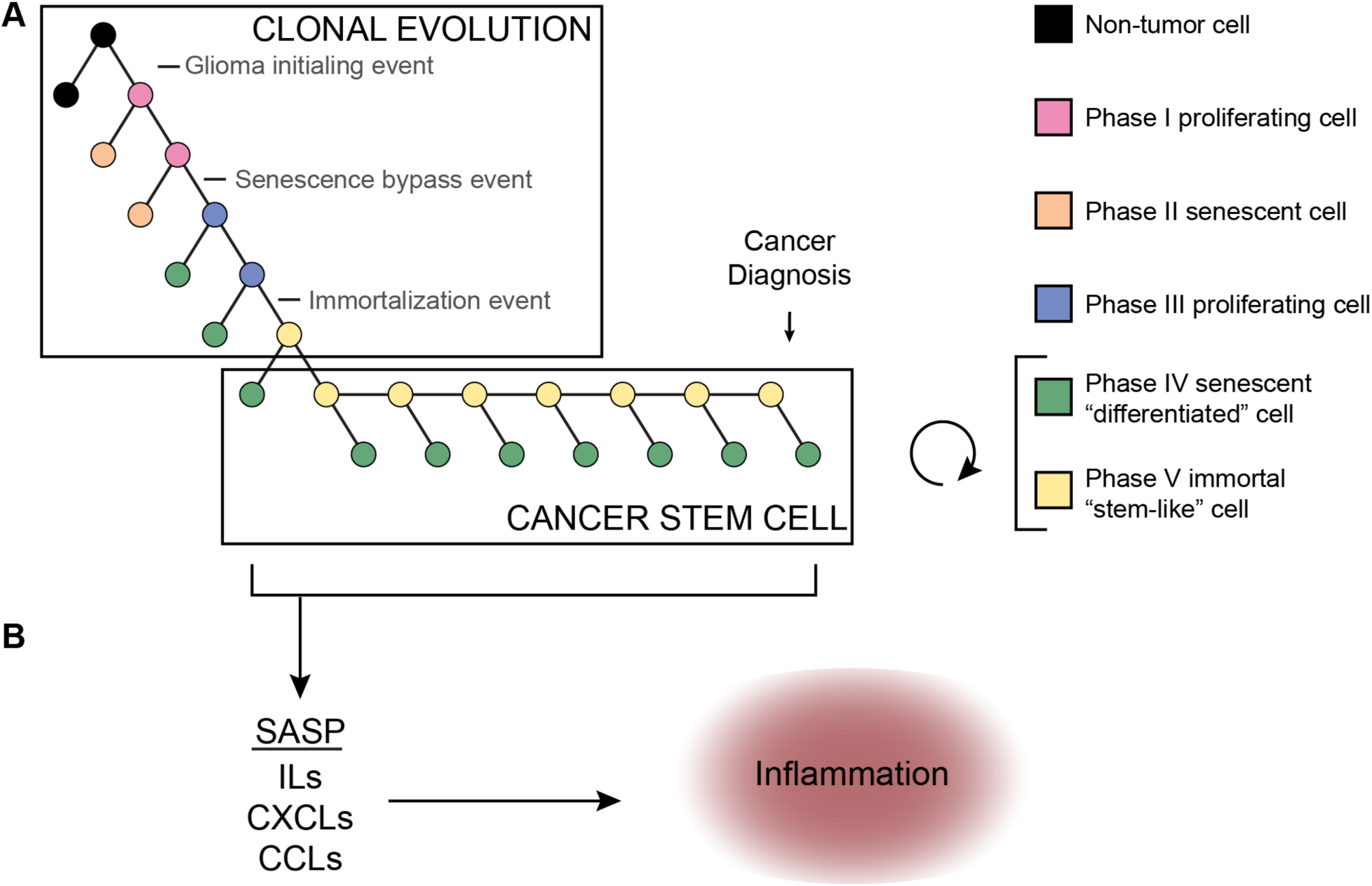
**A**. An integrated clonal evolution/cancer stem cell model for gliomagenesis assumes that sequential mutations and selection pressure drive the evolution of cancer stem-like cells. At the same time these stem-like cells may give rise to more differentiated (ie. phase IV) offspring that may divide further but rapidly become growth arrested. **B.** According to our model these cells may be senescent and contribute to the cancer phenotype by eliciting a micro-environment response via SASP. SASP = senescence-associated secretory phenotype; ILs = interleukines; CXCLs = chemokines (C-X-C motif); CCLs = chemokines (C-C motif)

Aforementioned studies supporting the cancer stem cell hypothesis suggest that multiple cellular populations exist within a tumor, including a self-renewing cancer stem cell population and a less-proliferative differentiated population. A recent paper demonstrated that cells derived from glioma stem cells may differentiate and subsequently undergo senescence [198]. We speculate that phase V tumor cells represent cancer stem cells that may give rise to more differentiated, phase IV cells (Figure 5A). The transition from phase IV to V and vice versa is likely volatile in nature owing to transcriptional reprogramming including the activation of the stemness factors Oligodendrocyte Transcription Factor 2 (*OLIG2*), Sex Determining Region Y – Box 2 (*SOX2)* and the reactivation of telomerase [199]. On the other hand, the transition from phase I to phase IV is more rigid in nature, involving various changes on a genomic level, including somatic mutations and copy number changes as described. This important distinction in flexibility led us to believe that phase V cells may re-differentiate and assume a phase IV state.

Although they are growth arrested, senescent cells are metabolically active and release a plethora of signaling molecules to the microenvironment. The senescence associated secretory phenotype (SASP) is a feature of senescent cells that curtails these cells with the release of proinflammatory cytokines [200]. SASP does not depend on p16^INK^4^a^ or p21 activity and senescence with intact p16^INKa^ function actually suppresses SASP [201]. Similarly, activated p53 signaling also suppresses SASP while *TP53* loss induces SASP [202-204]. These findings suggest that the SASP response is stronger when senescence pathway genes are lost. Indeed, IDH-wildtype astrocytomas often harbor homozygous deletions in *CDKN2A/B* and are known to have a highly active microenvironment [205]. Genes associated with SASP were shown to be overexpressed in higher grades of glioma and older patients, the latter group more likely to be affected by high-grade IDH-wildtype astrocytoma [206]. Moreover, primary glioblastoma cells retain a functional senescence program despite mutations in the *TERT* promoter and *CDKN2A/B* locus [207]. These findings imply that senescent and differentiated phase IV cells may be crucial for shaping the immune microenvironment in gliomas (Figure 5B).

## 4 A Broader Perspective

While there are not many known environmental risk factors that predispose to glioma, large cohort genome wide association studies over the past two decades have identified multiple heritable polymorphisms conferring glioma risk [35, 208-212]. Notably, several of these risk loci are localized to genes involved in telomere maintenance, including the telomerase reverse transcriptase *TERT*, telomerase RNA component *TERC*, and other telomere maintenance associated genes STN1, CST Complex Subunit (*OBFC1),* Protection Of Telomeres 1 (*POT1*) and Regulator of Telomere Elongation 1 (*RTEL1*) [208]. Moreover, there appears to be a significant increased glioma risk in people with increased leukocyte telomere lengths [213]. Unsurprisingly, glioma risk alleles at aforementioned genes are also associated with increased leukocyte telomere length [214-216]. Telomeres thus play an important role in not only the development of gliomas, but also in glioma risk [217]. In fact, the positive association between leukocyte telomere length and cancer risk is not specific to glioma and shared across many cancers. A recent Mendelian randomization study found that longer leukocyte telomere length was associated with an increased risk to cancer but a reduced risk to non-neoplastic disease such as cardiovascular disease [218]. It has been suggested that this difference is due to individuals with longer telomeres being more likely to acquire driver mutations due to an increased proliferative potential whereas the inverse relationship observed for non-neoplastic disease may be due to the impact of telomere shortening on tissue degeneration [219, 220].

Several hereditary disorders are associated with an increased risk of glioma development, including neurofibromatosis and the *TP53* germline mutation/Li-Fraumeni syndrome. Neurofibromatosis type I and II are autosomal dominant hereditary disorders with germline mutations in *NF1* and *NF2* and clinically characterized by multiple (non-gliomatous) benign nerve sheath tumors (especially neurofibromas in NF1, schwannomas in NF2), but also by a markedly increased risk on particular gliomas (especially pilocytic astrocytoma in NF1 and ependymomas in NF2) *[221-223]*. Both genes are well-known tumor suppressor genes and key components in the MAPK-pathway [224]. It has been demonstrated that senescence commonly occurs in benign nerve sheath tumors and that prolonged *NF1* disruption leads to oncogene-induced senescence in a model system, providing a rationale as to why these germline disorders present with tumors that are often relatively indolent [225]. A germline perturbation affecting *NF1* or *NF2* can be considered a tumor initiating event, explaining why this germline disorder guarantees the formation of multiple benign nerve sheath tumors over one’s lifetime.

Li-Fraumeni syndrome is a rare autosomal dominant hereditary disorder that is caused by the germline perturbation of *TP53* or *CHK2*, which regulates p53 activity [226-228]. Whereas neurofibromatosis guarantees the formation of multiple benign tumors in a lifetime, Li-Fraumeni patients pose a greater risk to developing a malignant tumor, e.g. a glioma [229]. This risk increases with age and is over 50% at age 30, with a lifetime cancer risk of over 70% in men and almost 100% in women [227]. Moreover, 15% and 4% of affected individuals were found to develop a second and third cancer [230]. Li-Fraumeni syndrome germline mutations affect phase II and prevent onset of oncogene-induced senescence following the acquisition of a glioma initiating event, thus increasing the risk of developing cancer over a lifetime.

The germline mutations underlying NF1/NF2 and Li-Fraumeni syndrome represent pathways that both need to be disrupted for a malignant tumor to form. The fact that nearly all patients with NF1/NF2 develop one or more benign tumors can be understood by acknowledging that in these disorders a germline tumor initiating event is involved. Unless this pathway is supplemented by a senescence bypass event these tumors do not readily proceed to malignancy. In contrast, Li-Fraumeni syndrome is characterized by an increased risk for malignant tumors in many (but not all) patients with this syndrome. In this syndrome, the germline senescence bypass event needs to be supplemented by a tumor initiating event for a tumor to develop, and such tumors may be more aggressive/malignant due to the defective senescence barrier, allowing the tumor (precursor) cell to instantly progress to phase III and instigate genomic instability.

## 5 Conclusion

Our knowledge on the molecular events driving cancer has grown exponentially over the years. This review has aimed to put this new knowledge into the perspective of the temporal molecular pathogenesis of glioma, starting from the first aberrant cell all the way to a symptom-causing glioma. To this end we have combined what is known on gene (mal)function, tumor evolution, genomic instability and telomere maintenance to develop a model of gliomagenesis. This model describes five sequential phases cancer cells undergo during their gliomagenesis. We speculate that transitions from one phase to the next can be characterized by acquisition of tumor driving events that sequentially contribute to the hallmarks of cancer as previously proposed, including proliferation, evasion of apoptosis and limitless replicative potential [231, 232]. Our model is a simplified abstraction of what may be the truth and new insights will refine and improve our understanding. Meanwhile, we hope that our model will help foster hypotheses leading to new insights on the molecular life history of glioma, that will help identify convincing therapeutic vulnerabilities.

## ACKNOWLEDGEMENTS

We thank members of the Verhaak laboratory for insightful discussion and valuable input. We thank Zoë Reifsnyder of the Jackson Laboratory creative team for artwork. This work was supported by grants from the National Institutes of Health R01 CA190121 and P30CA034196, the National Brain Tumor Society Oligo Research Fund and DefeatGBM Initiative, and grant 11026 from the Dutch Cancer Society KWF. We have no disclosures to make.

## REFERENCES

1. Vogelstein, B., et al., Cancer genome landscapes. Science, 2013. 339(6127): p. 1546–58.

2. Tomasetti, C., et al., Only three driver gene mutations are required for the development of lung and colorectal cancers. Proc Natl Acad Sci U S A, 2015. 112(1): p. 118–23.

3. Vogelstein, B. and K.W. Kinzler, The Path to Cancer—Three Strikes and You’re Out. N Engl J Med, 2015. 373(20): p. 1895–8.

4. Cairncross, G., et al., Phase III trial of chemoradiotherapy for anaplastic oligodendroglioma: long-term results of RTOG 9402. J Clin Oncol, 2013. 31(3): p. 337–43.

5. Cairncross, J.G., et al., Specific genetic predictors of chemotherapeutic response and survival in patients with anaplastic oligodendrogliomas. J Natl Cancer Inst, 1998. 90(19): p. 1473–9.

6. Wright, W.E., O.M. Pereira-Smith, and J.W. Shay, Reversible cellular senescence: implications for immortalization of normal human diploid fibroblasts. Mol Cell Biol, 1989. 9(7): p. 3088–92.

7. Romanov, S.R., et al., Normal human mammary epithelial cells spontaneously escape senescence and acquire genomic changes. Nature, 2001. 409(6820): p. 633–7.

8. Hahn, W.C., et al., Creation of human tumour cells with defined genetic elements. Nature, 1999. 400(6743): p. 464–8.

9. Ostrom, Q.T., et al., CBTRUS Statistical Report: Primary Brain and Central Nervous System Tumors Diagnosed in the United States in 2008-2012. Neuro Oncol, 2015. 17 Suppl 4: p. iv1–iv62.

10. Perry, A. and P. Wesseling, Histologic classification of gliomas. Handb Clin Neurol, 2016. 134: p. 71–95.

11. van den Bent, M.J., Interobserver variation of the histopathological diagnosis in clinical trials on glioma: a clinician’s perspective. Acta Neuropathol, 2010. 120(3): p. 297–304.

12. Coons, S.W., et al., Improving diagnostic accuracy and interobserver concordance in the classification and grading of primary gliomas. Cancer, 1997. 79(7): p. 1381–93.

13. Aldape, K., et al., Discrepancies in diagnoses of neuroepithelial neoplasms: the San Francisco Bay Area Adult Glioma Study. Cancer, 2000. 88(10): p. 2342–9.

14. Eckel-Passow, J.E., et al., Glioma Groups Based on 1p/19q, IDH, and TERT Promoter Mutations in Tumors. N Engl J Med, 2015. 372(26): p. 2499–508.

15. Network, C.G.A.R., et al., Comprehensive, Integrative Genomic Analysis of Diffuse Lower-Grade Gliomas. N Engl J Med, 2015. 372(26): p. 2481–98.

16. Wiestler, B., et al., Assessing CpG island methylator phenotype, 1p/19q codeletion, and MGMT promoter methylation from epigenome-wide data in the biomarker cohort of the NOA-04 trial. Neuro Oncol, 2014.

17. Ceccarelli, M., et al., Molecular Profiling Reveals Biologically Discrete Subsets and Pathways of Progression in Diffuse Glioma. Cell, 2016. 164(3): p. 550–63.

18. Yan, H., et al., IDH1 and IDH2 mutations in gliomas. The New England journal of medicine, 2009. 360(8): p. 765–773.

19. Louis, D.N., D. Schiff, and T.T. Batchelor, Classification of gliomas, in In: UpToDate, J.S. Loeffler, P.Y. Wen, and A.F. Eichler, Editors. 2014, UpToDate, Waltham, MA: Retrieved from http://www.uptodate.com/home/index.html.

20. Louis, D.N., et al., International Society Of Neuropathology—Haarlem consensus guidelines for nervous system tumor classification and grading. Brain Pathol, 2014. 24(5): p. 429–35.

21. Louis, D.N., et al., The 2016 World Health Organization Classification of Tumors of the Central Nervous System: a summary. Acta Neuropathol, 2016. 131(6): p. 803–20.

22. Barresi, V., et al., Dual-Genotype Diffuse Low-Grade Glioma: Is It Really Time to Abandon Oligoastrocytoma As a Distinct Entity? J Neuropathol Exp Neurol, 2017. 76(5): p. 342–346.

23. Schwartzentruber, J., et al., Driver mutations in histone H3.3 and chromatin remodelling genes in paediatric glioblastoma. Nature, 2012. 482(7384): p. 226–31.

24. Wu, G., et al., The genomic landscape of diffuse intrinsic pontine glioma and pediatric non-brainstem high-grade glioma. Nat Genet, 2014. 46(5): p. 444–50.

25. Korshunov, A., et al., Integrated analysis of pediatric glioblastoma reveals a subset of biologically favorable tumors with associated molecular prognostic markers. Acta Neuropathol, 2015. 129(5): p. 669–78.

26. Schindler, G., et al., Analysis of BRAF V600E mutation in 1,320 nervous system tumors reveals high mutation frequencies in pleomorphic xanthoastrocytoma, ganglioglioma and extra-cerebellar pilocytic astrocytoma. Acta Neuropathol, 2011. 121(3): p. 397–405.

27. Kepes, J.J., L.J. Rubinstein, and L.F. Eng, Pleomorphic xanthoastrocytoma: a distinctive meningocerebral glioma of young subjects with relatively favorable prognosis. A study of 12 cases. Cancer, 1979. 44(5): p. 1839–52.

28. Collins, V.P., D.T. Jones, and C. Giannini, Pilocytic astrocytoma: pathology, molecular mechanisms and markers. Acta Neuropathol, 2015. 129(6): p. 775–88.

29. Olar, A. and K.D. Aldape, Using the molecular classification of glioblastoma to inform personalized treatment. J Pathol, 2014. 232(2): p. 165–77.

30. Reuss, D.E., et al., ATRX and IDH1-R132H immunohistochemistry with subsequent copy number analysis and IDH sequencing as a basis for an "integrated" diagnostic approach for adult astrocytoma, oligodendroglioma and glioblastoma. Acta Neuropathol, 2015. 129(1): p. 133–46.

31. Ashley, D.J., The two "hit" and multiple "hit" theories of carcinogenesis. Br J Cancer, 1969. 23(2): p. 313–28.

32. Nordling, C.O., A new theory on cancer-inducing mechanism. Br J Cancer, 1953. 7(1): p. 68–72.

33. Nowell, P.C., The clonal evolution of tumor cell populations. Science, 1976. 194(4260): p. 23–8.

34. Knudson, A.G., Jr., Mutation and cancer: statistical study of retinoblastoma. Proc Natl Acad Sci U S A, 1971. 68(4): p. 820–3.

35. Shete, S., et al., Genome-wide association study identifies five susceptibility loci for glioma. Nat Genet, 2009. 41(8): p. 899–904.

36. Flavahan, W.A., E. Gaskell, and B.E. Bernstein, Epigenetic plasticity and the hallmarks of cancer. Science, 2017. 357(6348): p. eaal2380.

37. McGranahan, N. and C. Swanton, Clonal Heterogeneity and Tumor Evolution: Past, Present, and the Future. Cell, 2017. 168(4): p. 613–628.

38. Yates, L.R. and P.J. Campbell, Evolution of the cancer genome. Nat Rev Genet, 2012. 13(11): p. 795–806.

39. Johnson, B.E., et al., Mutational analysis reveals the origin and therapy-driven evolution of recurrent glioma. Science, 2014. 343(6167): p. 189–93.

40. Suzuki, H., et al., Mutational landscape and clonal architecture in grade II and III gliomas. Nat Genet, 2015. 47(5): p. 458–68.

41. Bai, H., et al., Integrated genomic characterization of IDH1-mutant glioma malignant progression. Nat Genet, 2016. 48(1): p. 59–66.

42. Suzuki, H., et al., Mutational landscape and clonal architecture in grade II and III gliomas. Nat Genet, 2015.

43. Lee, J.K., et al., Spatiotemporal genomic architecture informs precision oncology in glioblastoma. Nat Genet, 2017. 49(4): p. 594–599.

44. Kim, H., et al., Whole-genome and multisector exome sequencing of primary and post-treatment glioblastoma reveals patterns of tumor evolution. Genome Res, 2015. 25(3): p. 316–27.

45. Lu, C., et al., IDH mutation impairs histone demethylation and results in a block to cell differentiation. Nature, 2012. 483(7390): p. 474–478.

46. Turcan, S., et al., IDH1 mutation is sufficient to establish the glioma hypermethylator phenotype. Nature, 2012. 483(7390): p. 479–483.

47. McCarthy, N., Metabolism: unmasking an oncometabolite. Nat Rev Cancer, 2012. 12(4): p. 229.

48. Prensner, J.R. and A.M. Chinnaiyan, Metabolism unhinged: IDH mutations in cancer. Nat Med, 2011. 17(3): p. 291–3.

49. Koivunen, P., et al., Transformation by the (R)-enantiomer of 2-hydroxyglutarate linked to EGLN activation. Nature, 2012. 483(7390): p. 484–8.

50. Ward, P.S., et al., The common feature of leukemia-associated IDH1 and IDH2 mutations is a neomorphic enzyme activity converting alpha-ketoglutarate to 2-hydroxyglutarate. Cancer Cell, 2010. 17(3): p. 225–34.

51. Dang, L., et al., Cancer-associated IDH1 mutations produce 2-hydroxyglutarate. Nature, 2009. 462(7274): p. 739–44.

52. Noushmehr, H., et al., Identification of a CpG island methylator phenotype that defines a distinct subgroup of glioma. Cancer Cell, 2010. 17(5): p. 510–22.

53. Xu, W., et al., Oncometabolite 2-hydroxyglutarate is a competitive inhibitor of alpha-ketoglutarate-dependent dioxygenases. Cancer Cell, 2011. 19(1): p. 17–30.

54. Figueroa, M.E., et al., Leukemic IDH1 and IDH2 mutations result in a hypermethylation phenotype, disrupt TET2 function, and impair hematopoietic differentiation. Cancer Cell, 2010. 18(6): p. 553–67.

55. Flavahan, W.A., et al., Insulator dysfunction and oncogene activation in IDH mutant gliomas. Nature, 2016. 529(7584): p. 110–4.

56. Bigner, S.H., et al., Specific chromosomal abnormalities in malignant human gliomas. Cancer Res, 1988. 48(2): p. 405–11.

57. Ichimura, K., et al., Distinct patterns of deletion on 10p and 10q suggest involvement of multiple tumor suppressor genes in the development of astrocytic gliomas of different malignancy grades. Genes Chromosomes Cancer, 1998. 22(1): p. 9–15.

58. Sottoriva, A., et al., Intratumor heterogeneity in human glioblastoma reflects cancer evolutionary dynamics. Proc Natl Acad Sci U S A, 2013. 110(10): p. 4009–14.

59. Ozawa, T., et al., Most human non-GCIMP glioblastoma subtypes evolve from a common proneural-like precursor glioma. Cancer Cell, 2014. 26(2): p. 288–300.

60. Wang, J., et al., Clonal evolution of glioblastoma under therapy. Nat Genet, 2016. 48(7): p. 768–76.

61. Gerstung, M., et al., The evolutionary history of 2,658 cancers. bioRxiv, 2017.

62. Killela, P.J., et al., TERT promoter mutations occur frequently in gliomas and a subset of tumors derived from cells with low rates of self-renewal. Proc Natl Acad Sci U S A, 2013. 110(15): p. 6021–6.

63. Solimini, N.L., et al., Recurrent hemizygous deletions in cancers may optimize proliferative potential. Science, 2012. 337(6090): p. 104–9.

64. Martini, M., et al., PI3K/AKT signaling pathway and cancer: an updated review. Ann Med, 2014. 46(6): p. 372–83.

65. Li, J., et al., PTEN, a putative protein tyrosine phosphatase gene mutated in human brain, breast, and prostate cancer. Science, 1997. 275(5308): p. 1943–7.

66. Steck, P.A., et al., Identification of a candidate tumour suppressor gene, MMAC1, at chromosome 10q23.3 that is mutated in multiple advanced cancers. Nat Genet, 1997. 15(4): p. 356–62.

67. Pennisi, E., New tumor suppressor found—twice. Science, 1997. 275(5308): p. 1876–8.

68. Verhaak, R.G., et al., Integrated genomic analysis identifies clinically relevant subtypes of glioblastoma characterized by abnormalities in PDGFRA, IDH1, EGFR, and NF1. Cancer Cell, 2010. 17(1): p. 98–110.

69. Sturm, D., et al., Hotspot mutations in H3F3A and IDH1 define distinct epigenetic and biological subgroups of glioblastoma. Cancer Cell, 2012. 22(4): p. 425–37.

70. Wu, G., et al., Somatic histone H3 alterations in pediatric diffuse intrinsic pontine gliomas and non-brainstem glioblastomas. Nat Genet, 2012. 44(3): p. 251–3.

71. Sturm, D., et al., Paediatric and adult glioblastoma: multiform (epi)genomic culprits emerge. Nat Rev Cancer, 2014. 14(2): p. 92–107.

72. Korshunov, A., et al., Histologically distinct neuroepithelial tumors with histone 3 G34 mutation are molecularly similar and comprise a single nosologic entity. Acta Neuropathol, 2016. 131(1): p. 137–46.

73. Bender, S., et al., Reduced H3K27me3 and DNA hypomethylation are major drivers of gene expression in K27M mutant pediatric high-grade gliomas. Cancer Cell, 2013. 24(5): p. 660–72.

74. Lewis, P.W., et al., Inhibition of PRC2 activity by a gain-of-function H3 mutation found in pediatric glioblastoma. Science, 2013. 340(6134): p. 857–61.

75. Castel, D., et al., Histone H3F3A and HIST1H3B K27M mutations define two subgroups of diffuse intrinsic pontine gliomas with different prognosis and phenotypes. Acta Neuropathol, 2015. 130(6): p. 815–27.

76. Cantwell-Dorris, E.R., J.J. O’Leary, and O.M. Sheils, BRAFV600E: implications for carcinogenesis and molecular therapy. Mol Cancer Ther, 2011. 10(3): p. 385–94.

77. Davies, H., et al., Mutations of the BRAF gene in human cancer. Nature, 2002. 417(6892): p. 949–54.

78. Jones, D.T., et al., Tandem duplication producing a novel oncogenic BRAF fusion gene defines the majority of pilocytic astrocytomas. Cancer Res, 2008. 68(21): p. 8673–7.

79. Forshew, T., et al., Activation of the ERK/MAPK pathway: a signature genetic defect in posterior fossa pilocytic astrocytomas. J Pathol, 2009. 218(2): p. 172–81.

80. Sanoudou, D., et al., Analysis of pilocytic astrocytoma by comparative genomic hybridization. Br J Cancer, 2000. 82(6): p. 1218–22.

81. Collado, M. and M. Serrano, Senescence in tumours: evidence from mice and humans. Nat Rev Cancer, 2010. 10(1): p. 51–7.

82. Hayflick, L. and P.S. Moorhead, The serial cultivation of human diploid cell strains. Exp Cell Res, 1961. 25: p. 585–621.

83. He, S. and N.E. Sharpless, Senescence in Health and Disease. Cell, 2017. 169(6): p. 1000–1011.

84. Halazonetis, T.D., V.G. Gorgoulis, and J. Bartek, An oncogene-induced DNA damage model for cancer development. Science, 2008. 319(5868): p. 1352–5.

85. Collado, M., M.A. Blasco, and M. Serrano, Cellular senescence in cancer and aging. Cell, 2007. 130(2): p. 223–33.

86. Sharpless, N.E. and C.J. Sherr, Forging a signature of in vivo senescence. Nat Rev Cancer, 2015. 15(7): p. 397–408.

87. Campisi, J. and F. d’Adda di Fagagna, Cellular senescence: when bad things happen to good cells. Nat Rev Mol Cell Biol, 2007. 8(9): p. 729–40.

88. Perez-Mancera, P.A., A.R. Young, and M. Narita, Inside and out: the activities of senescence in cancer. Nat Rev Cancer, 2014. 14(8): p. 547–58.

89. Shain, A.H., et al., The Genetic Evolution of Melanoma from Precursor Lesions. N Engl J Med, 2015. 373(20): p. 1926–36.

90. Meeker, A.K., et al., Telomere length abnormalities occur early in the initiation of epithelial carcinogenesis. Clin Cancer Res, 2004. 10(10): p. 3317–26.

91. Kuilman, T., et al., Oncogene-induced senescence relayed by an interleukin-dependent inflammatory network. Cell, 2008. 133(6): p. 1019–31.

92. Gray-Schopfer, V.C., et al., Cellular senescence in naevi and immortalisation in melanoma: a role for p16? Br J Cancer, 2006. 95(4): p. 496–505.

93. Michaloglou, C., et al., BRAFE600-associated senescence-like cell cycle arrest of human naevi. Nature, 2005. 436(7051): p. 720–4.

94. Shay, J.W., O.M. Pereira-Smith, and W.E. Wright, A role for both RB and p53 in the regulation of human cellular senescence. Exp Cell Res, 1991. 196(1): p. 33–9.

95. Wright, W.E. and J.W. Shay, The two-stage mechanism controlling cellular senescence and immortalization. Exp Gerontol, 1992. 27(4): p. 383–9.

96. Ciriello, G., et al., Mutual exclusivity analysis identifies oncogenic network modules. Genome Res, 2012. 22(2): p. 398–406.

97. Ohgaki, H., et al., Genetic pathways to glioblastoma: a population-based study. Cancer Res, 2004. 64(19): p. 6892–9.

98. Watanabe, T., et al., IDH1 mutations are early events in the development of astrocytomas and oligodendrogliomas. Am J Pathol, 2009. 174(4): p. 1149–53.

99. Harris, S.L. and A.J. Levine, The p53 pathway: positive and negative feedback loops. Oncogene, 2005. 24(17): p. 2899–908.

100. Bromberg, J.E. and M.J. van den Bent, Oligodendrogliomas: molecular biology and treatment. Oncologist, 2009. 14(2): p. 155–63.

101. Jeuken, J.W., et al., The nature and timing of specific copy number changes in the course of molecular progression in diffuse gliomas: further elucidation of their genetic “life story”. Brain Pathol, 2011. 21(3): p. 308–20.

102. van Thuijl, H.F., et al., Spatial and temporal evolution of distal 10q deletion, a prognostically unfavorable event in diffuse low-grade gliomas. Genome Biol, 2014. 15(9): p. 471.

103. Aihara, K., et al., Genetic and epigenetic stability of oligodendrogliomas at recurrence. Acta Neuropathol Commun, 2017. 5(1): p. 18.

104. Bettegowda, C., et al., Mutations in CIC and FUBP1 contribute to human oligodendroglioma. Science, 2011. 333(6048): p. 1453–5.

105. Hu, X., et al., Multigene signature for predicting prognosis of patients with 1p19q co-deletion diffuse glioma. Neuro Oncol, 2017. 19(6): p. 786–795.

106. Bagchi, A. and A.A. Mills, The quest for the 1p36 tumor suppressor. Cancer Res, 2008. 68(8): p. 2551–6.

107. Vestin, A. and A.A. Mills, The tumor suppressor Chd5 is induced during neuronal differentiation in the developing mouse brain. Gene Expr Patterns, 2013. 13(8): p. 482–9.

108. Bagchi, A., et al., CHD5 is a tumor suppressor at human 1p36. Cell, 2007. 128(3): p. 459–75.

109. Kaghad, M., et al., Monoallelically expressed gene related to p53 at 1p36, a region frequently deleted in neuroblastoma and other human cancers. Cell, 1997. 90(4): p. 809–19.

110. Liggett, W.H., Jr. and D. Sidransky, Role of the p16 tumor suppressor gene in cancer. J Clin Oncol, 1998. 16(3): p. 1197–206.

111. Uhrbom, L., et al., Ink4a-Arf loss cooperates with KRas activation in astrocytes and neural progenitors to generate glioblastomas of various morphologies depending on activated Akt. Cancer Res, 2002. 62(19): p. 5551–8.

112. Holland, E.C., et al., Modeling mutations in the G1 arrest pathway in human gliomas: overexpression of CDK4 but not loss of INK4a-ARF induces hyperploidy in cultured mouse astrocytes. Genes Dev, 1998. 12(23): p. 3644–9.

113. Uhrbom, L., M. Nister, and B. Westermark, Induction of senescence in human malignant glioma cells by p16INK4A. Oncogene, 1997. 15(5): p. 505–14.

114. Ohgaki, H. and P. Kleihues, Genetic pathways to primary and secondary glioblastoma. Am J Pathol, 2007. 170(5): p. 1445–53.

115. Raabe, E.H., et al., BRAF activation induces transformation and then senescence in human neural stem cells: a pilocytic astrocytoma model. Clin Cancer Res, 2011. 17(11): p. 3590–9.

116. Horbinski, C., To BRAF or not to BRAF: is that even a question anymore? J Neuropathol Exp Neurol, 2013. 72(1): p. 2–7.

117. Koelsche, C., et al., Mutant BRAF V600E protein in ganglioglioma is predominantly expressed by neuronal tumor cells. Acta Neuropathol, 2013. 125(6): p. 891–900.

118. Zhang, J., et al., Whole-genome sequencing identifies genetic alterations in pediatric low-grade gliomas. Nat Genet, 2013. 45(6): p. 602–12.

119. Turcan, S. and T.A. Chan, MAPping the genomic landscape of low-grade pediatric gliomas. Nat Genet, 2013. 45(8): p. 847–9.

120. Jones, D.T., et al., Recurrent somatic alterations of FGFR1 and NTRK2 in pilocytic astrocytoma. Nat Genet, 2013. 45(8): p. 927–32.

121. Rozen, W.M., S. Joseph, and P.A. Lo, Spontaneous regression of low-grade gliomas in pediatric patients without neurofibromatosis. Pediatr Neurosurg, 2008. 44(4): p. 324–8.

122. Burkhard, C., et al., A population-based study of the incidence and survival rates in patients with pilocytic astrocytoma. J Neurosurg, 2003. 98(6): p. 1170–4.

123. Gunny, R.S., et al., Spontaneous regression of residual low-grade cerebellar pilocytic astrocytomas in children. Pediatr Radiol, 2005. 35(11): p. 1086–91.

124. Buder, T., et al., Model-Based Evaluation of Spontaneous Tumor Regression in Pilocytic Astrocytoma. PLoS Comput Biol, 2015. 11(12): p. e1004662.

125. Robinson, J.P., et al., Activated BRAF induces gliomas in mice when combined with Ink4a/Arf loss or Akt activation. Oncogene, 2010. 29(3): p. 335–44.

126. Huillard, E., et al., Cooperative interactions of BRAFV600E kinase and CDKN2A locus deficiency in pediatric malignant astrocytoma as a basis for rational therapy. Proc Natl Acad Sci U S A, 2012. 109(22): p. 8710–5.

127. Schiffman, J.D., et al., Oncogenic BRAF mutation with CDKN2A inactivation is characteristic of a subset of pediatric malignant astrocytomas. Cancer Res, 2010. 70(2): p. 512–9.

128. O’Sullivan, R.J. and J. Karlseder, Telomeres: protecting chromosomes against genome instability. Nat Rev Mol Cell Biol, 2010. 11(3): p. 171–81.

129. Olovnikov, A.M., A theory of marginotomy. The incomplete copying of template margin in enzymic synthesis of polynucleotides and biological significance of the phenomenon. J Theor Biol, 1973. 41(1): p. 181–90.

130. de Lange, T., How telomeres solve the end-protection problem. Science, 2009. 326(5955): p. 948–52.

131. Maser, R.S. and R.A. DePinho, Connecting chromosomes, crisis, and cancer. Science, 2002. 297(5581): p. 565–9.

132. McClintock, B., The Production of Homozygous Deficient Tissues with Mutant Characteristics by Means of the Aberrant Mitotic Behavior of Ring-Shaped Chromosomes. Genetics, 1938. 23(4): p. 315–76.

133. McClintock, B., The Stability of Broken Ends of Chromosomes in Zea Mays. Genetics, 1941. 26(2): p. 234–82.

134. Gisselsson, D., et al., Chromosomal breakage-fusion-bridge events cause genetic intratumor heterogeneity. Proc Natl Acad Sci U S A, 2000. 97(10): p. 5357–62.

135. Artandi, S.E., et al., Telomere dysfunction promotes non-reciprocal translocations and epithelial cancers in mice. Nature, 2000. 406(6796): p. 641–5.

136. Maciejowski, J., et al., Chromothripsis and Kataegis Induced by Telomere Crisis. Cell, 2015. 163(7): p. 1641–54.

137. Ernst, A., et al., Telomere dysfunction and chromothripsis. Int J Cancer, 2016. 138(12): p. 2905–14.

138. Maciejowski, J. and T. de Lange, Telomeres in cancer: tumour suppression and genome instability. Nat Rev Mol Cell Biol, 2017. 18(3): p. 175–186.

139. Notta, F., et al., A renewed model of pancreatic cancer evolution based on genomic rearrangement patterns. Nature, 2016. 538(7625): p. 378–382.

140. deCarvalho, A.C., et al., Discordant inheritance of chromosomal and extrachromosomal DNA elements contributes to dynamic disease evolution in glioblastoma. bioRxiv, 2017.

141. Turner, K.M., et al., Extrachromosomal oncogene amplification drives tumour evolution and genetic heterogeneity. Nature, 2017. 543(7643): p. 122–125.

142. Zheng, S., et al., A survey of intragenic breakpoints in glioblastoma identifies a distinct subset associated with poor survival. Genes Dev, 2013. 27(13): p. 1462–72.

143. Cox, D., C. Yuncken, and A.I. Spriggs, MINUTE CHROMATIN BODIES IN MALIGNANT TUMOURS OF CHILDHOOD. Lancet, 1965. 1(7402): p. 55–8.

144. Kohl, N.E., et al., Transposition and amplification of oncogene-related sequences in human neuroblastomas. Cell, 1983. 35(2 Pt 1): p. 359–67.

145. Sanborn, J.Z., et al., Double minute chromosomes in glioblastoma multiforme are revealed by precise reconstruction of oncogenic amplicons. Cancer Res, 2013. 73(19): p. 6036–45.

146. Nikolaev, S., et al., Extrachromosomal driver mutations in glioblastoma and low-grade glioma. Nat Commun, 2014. 5: p. 5690.

147. Kanda, T., K.F. Sullivan, and G.M. Wahl, Histone-GFP fusion protein enables sensitive analysis of chromosome dynamics in living mammalian cells. Curr Biol, 1998. 8(7): p. 377–85.

148. Forment, J.V., A. Kaidi, and S.P. Jackson, Chromothripsis and cancer: causes and consequences of chromosome shattering. Nat Rev Cancer, 2012. 12(10): p. 663–70.

149. Tanaka, H. and M.C. Yao, Palindromic gene amplification—an evolutionarily conserved role for DNA inverted repeats in the genome. Nat Rev Cancer, 2009. 9(3): p. 216–24.

150. Mazor, T., et al., Clonal expansion and epigenetic reprogramming following deletion or amplification of mutant IDH1. Proc Natl Acad Sci U S A, 2017.

151. Luchman, H.A., et al., Spontaneous loss of heterozygosity leading to homozygous R132H in a patient-derived IDH1 mutant cell line. Neuro Oncol, 2013. 15(8): p. 979–80.

152. Johannessen, T.A., et al., Rapid Conversion of Mutant IDH1 from Driver to Passenger in a Model of Human Gliomagenesis. Mol Cancer Res, 2016. 14(10): p. 976–983.

153. Counter, C.M., et al., Telomere shortening associated with chromosome instability is arrested in immortal cells which express telomerase activity. Embo j, 1992. 11(5): p. 1921–9.

154. Garbe, J.C., et al., Immortalization of normal human mammary epithelial cells in two steps by direct targeting of senescence barriers does not require gross genomic alterations. Cell Cycle, 2014. 13(21): p. 3423–35.

155. Morales, C.P., et al., Absence of cancer-associated changes in human fibroblasts immortalized with telomerase. Nat Genet, 1999. 21(1): p. 115–8.

156. Kim, N.W., et al., Specific association of human telomerase activity with immortal cells and cancer. Science, 1994. 266(5193): p. 2011–5.

157. Greider, C.W. and E.H. Blackburn, Identification of a specific telomere terminal transferase activity in Tetrahymena extracts. Cell, 1985. 43(2 Pt 1): p. 405–13.

158. Shay, J.W. and S. Bacchetti, A survey of telomerase activity in human cancer. Eur J Cancer, 1997. 33(5): p. 787–91.

159. Bryan, T.M., et al., Evidence for an alternative mechanism for maintaining telomere length in human tumors and tumor-derived cell lines. Nat Med, 1997. 3(11): p. 1271–4.

160. Heaphy, C.M., et al., Prevalence of the alternative lengthening of telomeres telomere maintenance mechanism in human cancer subtypes. Am J Pathol, 2011. 179(4): p. 1608–15.

161. Dunham, M.A., et al., Telomere maintenance by recombination in human cells. Nat Genet, 2000. 26(4): p. 447–50.

162. Heaphy, C.M., et al., Altered telomeres in tumors with ATRX and DAXX mutations. Science, 2011. 333(6041): p. 425.

163. Jiao, Y., et al., Frequent ATRX, CIC, FUBP1 and IDH1 mutations refine the classification of malignant gliomas. Oncotarget, 2012. 3(7): p. 709–22.

164. Kannan, K., et al., Whole-exome sequencing identifies ATRX mutation as a key molecular determinant in lower-grade glioma. Oncotarget, 2012. 3(10): p. 1194–203.

165. Goldberg, A.D., et al., Distinct factors control histone variant H3.3 localization at specific genomic regions. Cell, 2010. 140(5): p. 678–91.

166. Parkinson, G.N., M.P. Lee, and S. Neidle, Crystal structure of parallel quadruplexes from human telomeric DNA. Nature, 2002. 417(6891): p. 876–80.

167. Law, M.J., et al., ATR-X syndrome protein targets tandem repeats and influences allele-specific expression in a size-dependent manner. Cell, 2010. 143(3): p. 367–78.

168. Lazzerini-Denchi, E. and A. Sfeir, Stop pulling my strings - what telomeres taught us about the DNA damage response. Nat Rev Mol Cell Biol, 2016. 17(6): p. 364–78.

169. Lovejoy, C.A., et al., Loss of ATRX, genome instability, and an altered DNA damage response are hallmarks of the alternative lengthening of telomeres pathway. PLoS Genet, 2012. 8(7): p. e1002772.

170. Clynes, D., et al., Suppression of the alternative lengthening of telomere pathway by the chromatin remodelling factor ATRX. Nat Commun, 2015. 6: p. 7538.

171. Napier, C.E., et al., ATRX represses alternative lengthening of telomeres. Oncotarget, 2015. 6(18): p. 16543–58.

172. Ramamoorthy, M. and S. Smith, Loss of ATRX Suppresses Resolution of Telomere Cohesion to Control Recombination in ALT Cancer Cells. Cancer Cell, 2015. 28(3): p. 357–69.

173. Horn, S., et al., TERT promoter mutations in familial and sporadic melanoma. Science, 2013. 339(6122): p. 959–61.

174. Vinagre, J., et al., Frequency of TERT promoter mutations in human cancers. Nat Commun, 2013. 4: p. 2185.

175. Huang, F.W., et al., Highly recurrent TERT promoter mutations in human melanoma. Science, 2013. 339(6122): p. 957–9.

176. Bell, R.J.A., et al., The transcription factor GABP selectively binds and activates the mutant TERT promoter in cancer. Science (New York, N.Y.), 2015. 348(6238): p. 1036–1039.

177. Martinez, P. and M.A. Blasco, Telomeric and extra-telomeric roles for telomerase and the telomere-binding proteins. Nat Rev Cancer, 2011. 11(3): p. 161–76.

178. Park, J.I., et al., Telomerase modulates Wnt signalling by association with target gene chromatin. Nature, 2009. 460(7251): p. 66–72.

179. Chiba, K., et al., Cancer-associated TERT promoter mutations abrogate telomerase silencing. Elife, 2015. 4.

180. Chiba, K., et al., Mutations in the promoter of the telomerase gene TERT contribute to tumorigenesis by a two-step mechanism. Science, 2017.

181. Maryoung, L., et al., Somatic mutations in telomerase promoter counterbalance germline loss-of-function mutations. J Clin Invest, 2017. 127(3): p. 982–986.

182. Langford, L.A., et al., Telomerase activity in human brain tumours. Lancet, 1995. 346(8985): p. 1267–8.

183. Pekmezci, M., et al., Adult infiltrating gliomas with WHO 2016 integrated diagnosis: additional prognostic roles of ATRX and TERT. Acta Neuropathol, 2017. 133(6): p. 1001–1016.

184. Barthel, F.P., et al., Systematic analysis of telomere length and somatic alterations in 31 cancer types. Nat Genet, 2017. 49(3): p. 349–357.

185. Mangerel, J., et al., Alternative lengthening of telomeres is enriched in, and impacts survival of TP53 mutant pediatric malignant brain tumors. Acta Neuropathol, 2014. 128(6): p. 853–62.

186. Lee, J., D.A. Solomon, and T. Tihan, The role of histone modifications and telomere alterations in the pathogenesis of diffuse gliomas in adults and children. J Neurooncol, 2017. 132(1): p. 1–11.

187. Vescovi, A.L., R. Galli, and B.A. Reynolds, Brain tumour stem cells. Nat Rev Cancer, 2006. 6(6): p. 425–36.

188. Greaves, M. and C.C. Maley, Clonal evolution in cancer. Nature, 2012. 481(7381): p. 306–13.

189. Patel, A.P., et al., Single-cell RNA-seq highlights intratumoral heterogeneity in primary glioblastoma. Science, 2014. 344(6190): p. 1396–401.

190. Tirosh, I., et al., Single-cell RNA-seq supports a developmental hierarchy in human oligodendroglioma. Nature, 2016. 539(7628): p. 309–313.

191. Venteicher, A.S., et al., Decoupling genetics, lineages, and microenvironment in IDH-mutant gliomas by single-cell RNA-seq. Science, 2017. 355(6332).

192. Lan, X., et al., Fate mapping of human glioblastoma reveals an invariant stem cell hierarchy. Nature, 2017.

193. Kimura, M., The neutral theory of molecular evolution: a review of recent evidence. Jpn J Genet, 1991. 66(4): p. 367–86.

194. Williams, M.J., et al., Identification of neutral tumor evolution across cancer types. Nat Genet, 2016. 48(3): p. 238–244.

195. Noorbakhsh, J. and J.H. Chuang, Uncertainties in tumor allele frequencies limit power to infer evolutionary pressures. Nat Genet, 2017. 49(9): p. 1288–1289.

196. Tarabichi, M., et al., Neutral tumor evolution? bioRxiv, 2017.

197. Williams, M.J., et al., Reply: Uncertainties in tumor allele frequencies limit power to infer evolutionary pressures. Nat Genet, 2017. 49(9): p. 1289–1291.

198. Ouchi, R., et al., Senescence from glioma stem cell differentiation promotes tumor growth. Biochem Biophys Res Commun, 2016. 470(2): p. 275–281.

199. Suva, M.L., et al., Reconstructing and reprogramming the tumor-propagating potential of glioblastoma stem-like cells. Cell, 2014. 157(3): p. 580–94.

200. Coppe, J.P., et al., The senescence-associated secretory phenotype: the dark side of tumor suppression. Annu Rev Pathol, 2010. 5: p. 99–118.

201. Coppe, J.P., et al., Tumor suppressor and aging biomarker p16(INK4a) induces cellular senescence without the associated inflammatory secretory phenotype. J Biol Chem, 2011. 286(42): p. 36396–403.

202. Coppe, J.P., et al., Senescence-associated secretory phenotypes reveal cell-nonautonomous functions of oncogenic RAS and the p53 tumor suppressor. PLoS Biol, 2008. 6(12): p. 2853–68.

203. Raj, N. and L.D. Attardi, Tumor suppression: p53 alters immune surveillance to restrain liver cancer. Curr Biol, 2013. 23(12): p. R527–30.

204. Lujambio, A., et al., Non-cell-autonomous tumor suppression by p53. Cell, 2013. 153(2): p. 449–60.

205. Wang, Q., et al., Tumor Evolution of Glioma-Intrinsic Gene Expression Subtypes Associates with Immunological Changes in the Microenvironment. Cancer Cell, 2017. 32(1): p. 42–56.e6.

206. Coppola, D., et al., Senescence-associated-gene signature identifies genes linked to age, prognosis, and progression of human gliomas. J Geriatr Oncol, 2014. 5(4): p. 389–99.

207. Kumar, R., et al., Induction of senescence in primary glioblastoma cells by serum and TGFbeta. Sci Rep, 2017. 7(1): p. 2156.

208. Melin, B.S., et al., Genome-wide association study of glioma subtypes identifies specific differences in genetic susceptibility to glioblastoma and non-glioblastoma tumors. Nat Genet, 2017. 49(5): p. 789–794.

209. Wrensch, M., et al., Variants in the CDKN2B and RTEL1 regions are associated with high-grade glioma susceptibility. Nat Genet, 2009. 41(8): p. 905–8.

210. Rajaraman, P., et al., Genome-wide association study of glioma and meta-analysis. Hum Genet, 2012. 131(12): p. 1877–88.

211. Jenkins, R.B., et al., A low-frequency variant at 8q24.21 is strongly associated with risk of oligodendroglial tumors and astrocytomas with IDH1 or IDH2 mutation. Nat Genet, 2012. 44(10): p. 1122–5.

212. Kinnersley, B., et al., Genome-wide association study identifies multiple susceptibility loci for glioma. Nat Commun, 2015. 6: p. 8559.

213. Connelly, J.M. and M.G. Malkin, Environmental risk factors for brain tumors. Curr Neurol Neurosci Rep, 2007. 7(3): p. 208–14.

214. Codd, V., et al., Identification of seven loci affecting mean telomere length and their association with disease. Nat Genet, 2013. 45(4): p. 422–7, 427e1-2.

215. Walsh, K.M., et al., Longer genotypically-estimated leukocyte telomere length is associated with increased adult glioma risk. Oncotarget, 2015. 6(40): p. 42468–77.

216. Walsh, K.M., et al., Telomere maintenance and the etiology of adult glioma. Neuro Oncol, 2015. 17(11): p. 1445–52.

217. Walsh, K.M., et al., Variants near TERT and TERC influencing telomere length are associated with high-grade glioma risk. Nat Genet, 2014. 46(7): p. 731–5.

218. Haycock, P.C., et al., Association Between Telomere Length and Risk of Cancer and Non-Neoplastic Diseases: A Mendelian Randomization Study. JAMA Oncol, 2017. 3(5): p. 636–651.

219. Blasco, M.A., Telomere length, stem cells and aging. Nat Chem Biol, 2007. 3(10): p. 640–9.

220. Stone, R.C., et al., Telomere Length and the Cancer-Atherosclerosis Trade-Off. PLoS Genet, 2016. 12(7): p. e1006144.

221. Gutmann, D.H., et al., The diagnostic evaluation and multidisciplinary management of neurofibromatosis 1 and neurofibromatosis 2. Jama, 1997. 278(1): p. 51–7.

222. Cawthon, R.M., et al., A major segment of the neurofibromatosis type 1 gene: cDNA sequence, genomic structure, and point mutations. Cell, 1990. 62(1): p. 193–201.

223. Rouleau, G.A., et al., Alteration in a new gene encoding a putative membrane-organizing protein causes neuro-fibromatosis type 2. Nature, 1993. 363(6429): p. 515–21.

224. Cichowski, K., et al., Mouse models of tumor development in neurofibromatosis type 1. Science, 1999. 286(5447): p. 2172–6.

225. Courtois-Cox, S., et al., A negative feedback signaling network underlies oncogene-induced senescence. Cancer Cell, 2006. 10(6): p. 459–72.

226. Bell, D.W., et al., Heterozygous germ line hCHK2 mutations in Li-Fraumeni syndrome. Science, 1999. 286(5449): p. 2528–31.

227. McBride, K.A., et al., Li-Fraumeni syndrome: cancer risk assessment and clinical management. Nat Rev Clin Oncol, 2014. 11(5): p. 260–71.

228. Birch, J.M., et al., Prevalence and diversity of constitutional mutations in the p53 gene among 21 Li-Fraumeni families. Cancer Res, 1994. 54(5): p. 1298–304.

229. Birch, J.M., et al., Relative frequency and morphology of cancers in carriers of germline TP53 mutations. Oncogene, 2001. 20(34): p. 4621–8.

230. Hisada, M., et al., Multiple primary cancers in families with Li-Fraumeni syndrome. J Natl Cancer Inst, 1998. 90(8): p. 606–11.

231. Hanahan, D. and R.A. Weinberg, The hallmarks of cancer. Cell, 2000. 100(1): p. 57–70.

232. Hanahan, D. and R.A. Weinberg, Hallmarks of cancer: the next generation. Cell, 2011. 144(5): p. 646–74.

